# Structures of the Cmr-β Complex Reveal the Regulation of the Immunity Mechanism of Type III-B CRISPR-Cas

**DOI:** 10.1101/2020.06.24.163345

**Authors:** Nicholas Sofos, Mingxia Feng, Stefano Stella, Tillmann Pape, Anders Fuglsang, Jinzhong Lin, Qihong Huang, Yingjun Li, Qunxin She, Guillermo Montoya

## Abstract

Cmr-β is a Type III-B CRISPR-Cas complex that upon target RNA recognition unleashes a multifaceted immune response against invading genetic elements, including ssDNA cleavage, cyclic oligoadenylate synthesis, and also a unique UA-specific ssRNA hydrolysis by the Cmr2 subunit. Here, we present the structure-function relationship of Cmr-β unveiling how binding of the target RNA regulates the Cmr2 activities. CryoEM analysis revealed the unique subunit architecture of Cmr-β and captured the complex in different conformational stages of the immune response, including the non-cognate and cognate target-RNA bound complexes. The binding of the target RNA induces a conformational change of Cmr2, which together with the complementation between the 5’-handle in the crRNA and the 3’-antitag of the target RNA, activate different configurations in a unique loop of the Cmr3 subunit, which acts as an allosteric sensor signaling the self vs. non-self recognition. These findings highlight the diverse defense strategies of Type III complexes.

## INTRODUCTION

Prokaryotes employ RNA-guided adaptive immune systems called CRISPR-Cas (<u>c</u>lustered <u>r</u>egularly <u>i</u>nterspaced <u>s</u>hort <u>p</u>alindromic <u>r</u>epeat-<u>C</u>RISPR <u>as</u>sociated) against mobile genetic elements (MGEs) such as viruses and plasmids (Barrangou et al., 2007; Makarova et al., 2006; Marraffini and Sontheimer, 2008). This immunity is acquired by integrating short DNA sequences (protospacers) derived from the invader into the host CRISPR locus (Amitai and Sorek, 2016). The CRISPR loci are transcribed and processed into CRISPR RNAs (crRNA) that contain a spacer typically flanked by parts of the repeat sequences on either side. The crRNA assembles with Cas proteins to form crRNA-guided ribonucleoprotein (crRNP) effector complexes that recognize and degrade foreign genetic elements in a crRNA-guided fashion (Hille et al., 2018). Based on the effector complex composition, CRISPR-Cas systems are divided into two classes and six types (Abudayyeh et al., 2016; Makarova et al., 2015).

Type III is of particular interest as they degrade both RNA and DNA of the invaders, and is subdivided into subtypes III-A to III-F with different subunits and architectures highlighting the large heterogeneity of this CRISPR-Cas system (Makarova et al., 2020). Subtype III-A (Csm) and III-B (Cmr) are the best characterized. Although they present differences in composition, both subtypes share the signature *cas10* gene product called Csm1 and Cmr2 respectively (Fig. S1A-B). The *cas10* gene encodes a multidomain protein containing an N-terminal HD (histidine–aspartate) domain followed by two Palm domains joined by a Zinc Finger (ZnF), a small α-helical domain (D2) and another C-terminal α-helical domain (D4) (Fig. S1A-B) (Mohanraju et al., 2016). Type III CRISPR-Cas systems deploy a unique multipronged approach to defend against foreign nucleic acid. As the host RNA polymerase transcribes the foreign DNA, the nascent mRNA is recognized by base pairing with complementary crRNA, and cleaved into ssRNA fragments at 6-nt intervals by the Cmr4/Csm3 subunits (Ramia et al., 2014; Staals et al., 2014; Tamulaitis et al., 2014). This binding tethers the complex to the transcription bubble (Amitai and Sorek, 2017; Kazlauskiene et al., 2017). Identification of the mRNA as an invader-derived transcript results in the activation of the Cmr2/Csm1 protein. The host transcript (self) is distinguished from an invader (non-self)-derived mRNA transcript based on the complementarity between an 8-nt repeat-derived crRNA 5’-tag, the 5’-repeat handle sequence (RHS), and the 3’-antitag of the target RNA, the 3’-protospacer flanking sequence (PFS). Complementarity between the 5’-tag and 3’-antitag of the non-cognate target RNA (NTR) triggers the effector complex self-inactivation, to prevent an autoimmune response (Elmore et al., 2016; Estrella et al., 2016; Kazlauskiene et al., 2016), while uncoupling between the 3’-antitag of an invader-derived transcript and the 5’-tag of the crRNA, cognate target RNA (CTR), triggers activation of the Cmr2/Csm1, resulting in ssDNA cleavage by the HD nuclease, and conversion of ATP into cyclic oligoadenylates (cOA) by the Palm domains of Cmr2/Csm1. cOAs act as secondary messengers by allosterically activating CRISPR-Cas type III associated RNases (Han et al., 2018; Kazlauskiene et al., 2017; Niewoehner et al., 2017) of the Csx1/Csm6 family for robust RNA decay (Jia et al., 2019b; Molina et al., 2019), leading to cell dormancy or death (Rouillon et al., 2018).

Recently, several studies of the Type III-A Csm complex have shown mechanistic details of NTR *vs.* CTR recognition, activation of ssDNA cleavage and cOA synthesis (Guo et al., 2019; Jia et al., 2019c; You et al., 2019). However, even though Csm and Cmr systems represent Type III, the A and B subtypes are considerably divergent in their sequences, as reflected in the different subunits and architectures. Currently, a full understanding of the Cmr complex and its activities is missing. The structural knowledge of Type III-B Cmr complexes is limited to a chimeric complex bound to a DNA analogue of the target RNA lacking the Cmr1 protein and Cmr2 HD domain (Osawa et al., 2015), and a medium resolution cryoEM reconstructions of *T. thermophilus* Cmr lacking the Cmr2 HD domain (Taylor et al., 2015). Here, we characterize *S. islandicus* (Sis) Cmr-β complex activities, and present seven high resolution cryoEM structures of the complete complex including the apo, NTR-, CTR-, and CTR-AMPPnP-bound states. Our findings show how SisCmr-β is able to discriminate between foreign and self mRNA transcripts, resulting in novel tactics of fine-tuning the immune response against invading MGEs. Furthermore, we report on a unique ability of SisCmr-β HD domain to cleave ssRNA specifically between U and A nucleotides, suggesting that this complex could also contribute to RNA decay during infection.

## RESULTS

### SisCmr-β cleaves ssDNA, generates cOA and additionally cuts non-complementary ssRNA

The genes coding for SisCmr-β occur at the same locus on the *S. islandicus* chromosome, while the gene coding for the SisCsx1 associated RNase is located next to the SisCmr-α locus (Fig. S2A). An additional subunit called Cmr7 has been identified in *S. islandicus* and *S. solfataricus* (Sso) Cmr-β and Cmr effector complexes (Deng et al., 2013; Zhang et al., 2012). We isolated SisCmr-β from *S. islandicus* (Han et al., 2018), which displayed a mass of ~900 kDa by MALS (Multiple Angle Light Scattering) (Fig. S2B-C, STAR Methods), indicating that SisCmr-β is larger than other Type III CRISPR-Cas complexes (Guo et al., 2019; Jia et al., 2019c; Osawa et al., 2015; You et al., 2019).

We performed enzymatic assays to characterize the biochemical activities of SisCmr-β (Fig. S2D-H). We first tested the cleavage of target RNA by SisCmr-β using three labeled target RNA molecules as a substrate (STAR Methods Table S1): CTR, with mismatches between its 3′-PFS and 5′-RHS of crRNA; naTR, a target RNA lacking the 3′-PFS; and NTR whose 3′-PFS was complementary to 5′-RHS of crRNA (Fig. S2D). In all these cases, cleavage of the target RNA resulted in the appearance of four fragments (17, 23, 29, and 35 oligonucleotides) at 6-nt length intervals (Fig. S2D-E). These experiments indicated that the target RNA can be cleaved independently of the hybridization status between the 5’-RHS and the 3’-PFS (Fig. S2E). We also monitored the ssDNA severing activity and cOA synthesis by the SisCmr-β complex (Fig. S2F–G). Unlike the target RNase activity, we observed ssDNA severing and cOA synthesis only upon CTR binding. This indicated that mismatches between the 5′-RHS of crRNA and the 3′-PFS of target RNA are essential for the activation of the *cas10* gene product, as previously observed (Elmore et al., 2016; Estrella et al., 2016; Han et al., 2017; Han et al., 2018; Kazlauskiene et al., 2017; Kazlauskiene et al., 2016; Niewoehner et al., 2017; Rouillon et al., 2018).

Next we asked whether the cOA moiety produced by SisCmr-β activates SisCsx1 as we previously demonstrated for cOA generated by SisCmr-α (Han et al., 2018; Molina et al., 2019). The reason for their co-existence is unclear. Both complexes generate cOAs, but SisCmr-α synthesizes approximately 10 times more cOA than SisCmr-β (Fig. S2I). The binding and activity assays revealed that cOA generated by SisCmr-β binds and activates SisCsx1 (Fig. S2J–K), supporting the notion that both complexes can activate the accessory Csx1 RNase. To identify important residues involved in SisCmr-β activities, we performed a sequence analysis including Cmr-α and Csm orthologues, which revealed the conservation of catalytic amino acids across different Type III effector complexes. This analysis indicated the key residues in SisCmr-β for specific target RNA degradation in the Cmr4 subunits (D31) (Osawa et al., 2015) as well as cOA formation (G800, G801, D803, D804, D650 and D652) and ssDNA degradation (H20 and D21) in Cmr2 (Fig. S1B). The identification of these key residues helped to reveal that SisCmr-β complex can cleave ssRNA that is not complementary to the crRNA. We observed additional cleavage sites on the target RNA when Cmr4 activity was abrogated by the D31A mutation (Fig. 2L). A discrete number of reaction products (bands), which matched the occurrence of U-A stretches in a ssRNA substrate (S10RNA) were observed upon addition of the CTR (Fig. S2D, S2H, STAR Methods Table S1), thus suggesting UA specificity of the SisCmr-β ssRNase. This activity is less efficient than unspecific ssDNA degradation. In addition, we observed that the Cmr2 H20A/D21A double variant, whose ssDNA-degrading activity was abrogated (Fig. S2M), also exhibited a severe impairment of the UA ssRNase activity (Fig. S2N). This indicated that the HD domain is involved in UA ssRNA severing. The target RNA cleavage and cOA synthesis were unaffected by the H20A/D21A variant (Fig. S2L, S2O). Collectively, these experiments supported the role of conserved amino acid residues in SisCmr-β activity and revealed the new UA ssRNase activity.

### Overall Structure of the SisCmr-β complex revealed by Cryo-EM

To investigate the coordination of the activities of this multienzymatic complex, we solved 7 different cryoEM structures of the apo, NTR and CTR target RNA-bound, as well as several CTR-bound states in the presence of AMP-PnP and a 20-nt polyT ssDNA, at global resolutions ranging from 2.4 Å to 3.5 Å (Fig. S3, S4). The apo structure of SisCmr-β at 2.75 Å global resolution (Fig. 1) Fig. S3, S4, STAR Methods Table S2), revealed the singular architecture of the complex, with crRNA in its core (Fig. 1A-B) and the protein subunits assembled around it, with a subunit stoichiometry of Cmr1_1_-2_1_-3_1_-4_4_-5_3_-6_1_-7_26_, including 13 dimers of the unique Cmr7 subunit decorating the crRNP particle (Fig. 1B-C). Previous studies revealed that Type III complexes combine two intertwined helical filaments of multiple stacked copies of Cmr4/Csm3 and Cmr5/Csm2 (Jia et al., 2019c; Osawa et al., 2015; You et al., 2019), wrapping around the crRNA and the crRNA-target RNA duplex in a Cmr4_n+1_Cmr5_n_ stoichiometry. We observed a similar helical filament in SisCmr-β, which assembles four Cmr4 and three Cmr5 subunits around the crRNA (Fig 1B-C), and was longer than that of any other Type III members.

**Figure 1.**
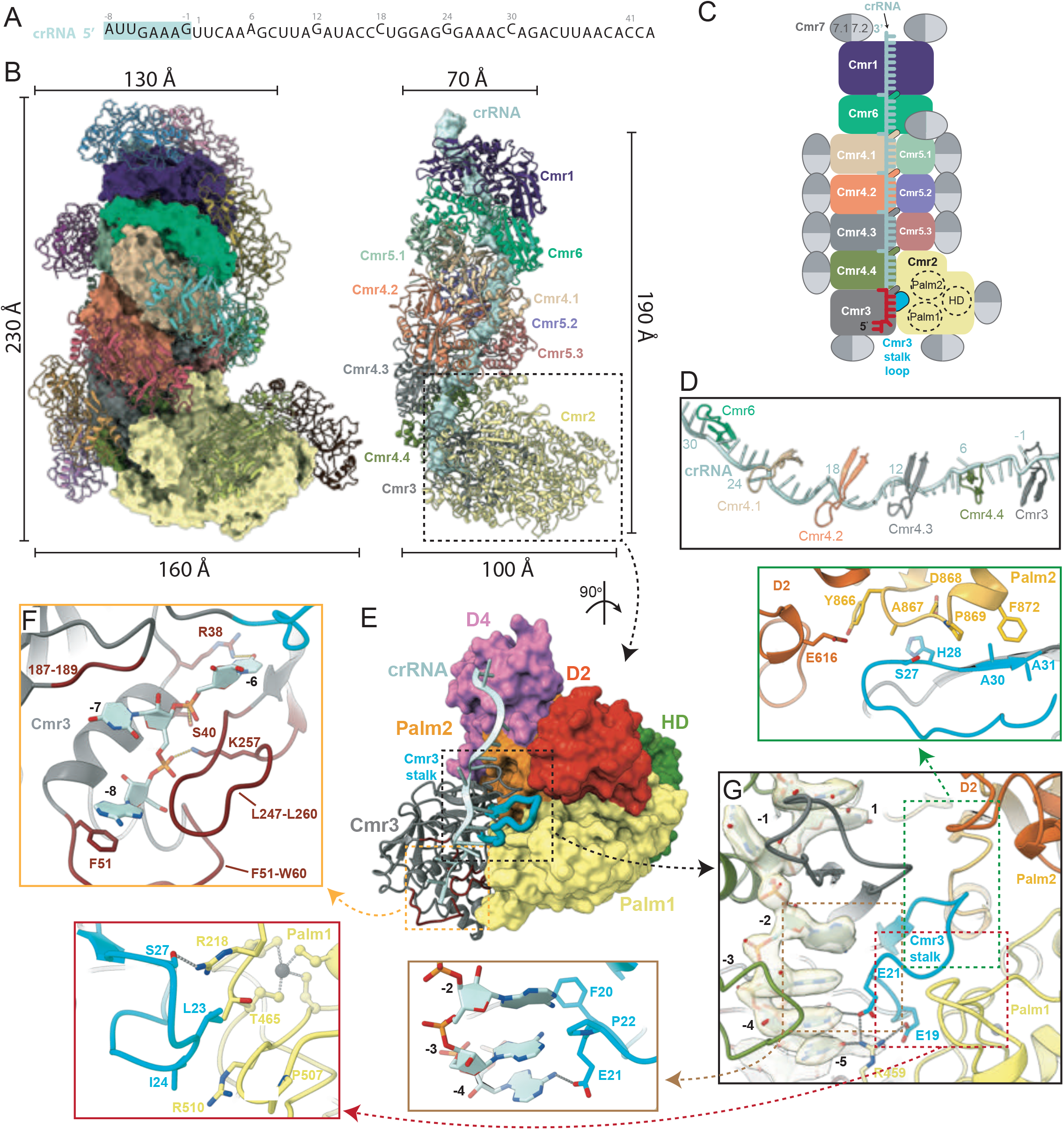
Overall Structure of the SisCmr-β Apo Complex Revealed by Cryo-EM. **(A)** Sequence of the crRNA as observed in the structure. The repeat-derived 5’-tag sequence, position −8 to −1, is shown in teal, followed by the spacer-derived region (position 1 to 43). **(B)** The overall SisCmr-β apo structure. Left, Cmr1-Cmr6 and the bound crRNA are shown in surface representation, and the Cmr7 dimers are in cartoon representation. Right, structure with the Cmr7 dimers removed, showcasing the inner filamentous core region formed by Cmr1-Cmr6 (cartoon representation), as well as the crRNA (surface representation). **(C)** Schematic representation of the SisCmr-β structure with the same color code used for Cmr1-Cmr6 as in 1B. The Cmr7 dimers are shown in light and dark grey, representing each subunit, and the binding sites of 13 Cmr7 dimers are indicated. The 5’-RHS is shown in red, and the Cmr3 stalk loop (residues 16-31) is shown in cyan. **(D)** The crRNA, and β-thumbs of the four Cmr4 subunits, Cmr3, and Cmr6. Each β-thumb folds over the crRNA, generating a kink in the crRNA and splaying out a base at 6-nt intervals. **(E)** Structure rotated ~90° degrees around the y-axis relative to the position shown in Fig. 1B, with the Cmr3 subunit is shown in grey cartoon with the stalk loop highlighted in cyan, bound to the crRNA and Cmr2 domain (surface representation). **(F)** Close-up of the crRNA 5’-tag region showing nucleotides −6 to −8 wrapped primarily by two loops, L51-60 and L247-260 of Cmr3. **(G)** Close-up of the −1 to −5 crRNA 5’-RHS region. Nucleotides –2 to –5 are exposed and able to engage in the self/non-self recognition of the 3′-PFS of bound target RNA. The density of crRNA is shown in surface representation. The Cmr3 stalk loop is wedged between nucleotides –2 to –5, the Cmr3 β-thumb, and Cmr2 Palm1, Palm2, and D2 domains. Arrows point to focused views showing detailed interactions of the Cmr3 stalk loop with the respective regions. See also Fig. S1 to S6.

The Cmr4 subunits contained a β-thumb, also observed in Csm3, which folds over the crRNA kinking its backbone (Fig. 1D). The RNase centers of the Cmr4 subunits, where the catalytic D31 residue resides, were similar to those of the Csm3 subunit of the Csm complex (Jia et al., 2019c; You et al., 2019) (Fig. S5A). However, there were some differences in the Cmr4 catalytic sites architecture; while Cmr4.1, 4.2, and 4.3 centers were assembled between the Cmr4 and Cmr5 subunits, the Cmr4.4 active site interacted with Cmr2 (Fig. 1C, Fig. S5B-D). In the apo structure of SisCmr-β complex, D31 in Cmr4.4 was associated with the strictly conserved Cmr2Y960 and Cmr2R1030 in the D4 domain, while D31 in Cmr4.3 was associated with the conserved Cmr5.3K145 and Cmr5.3R35 (Fig. S5B). By contrast, the catalytic D31 residues in Cmr4.2 and Cmr4.1 were not engaged in any interaction.

The large Cmr2 subunit, which contains the catalytic centers for cOA synthesis and ssDNA/UA ssRNA degradation, is positioned at the 5’-end of the crRNA together with Cmr3 (Fig. 1C, 1E). The crRNA 5′-RHS was attached to Cmr3 (Fig. 1F-G, Fig. S1C). The initial three nucleotides (A-8, U-7 and U-6) of the 5′-RHS were “wrapped” by two unique loops (L51–W60 and L247–L260) of Cmr3, making these nucleotides unavailable for coupling with the 3′-PFS of target RNA (Fig. 1F-G, Fig. S1C). A-8 was stacked with F51, while U-7 was stacked with the main chain of residues 187–189 of Cmr3. The phosphate backbone of A-8 to U-6 engaged in polar interactions with K257, I251, and S40, and U-6 was contacted by R38. Interestingly, these loops were stabilized by multiple contacts with the Palm1 and ZnF domains of Cmr2. As G-1 in the 5′-RHS was flipped out by the loop between S141–T153 of Cmr3 (Fig. 1G), only the G-5 to A-2 nucleotides were available for testing complementation with the target RNA 3′-PFS. In the apo structure, these nucleotides were protected by the arrangement promoted by a unique loop in Cmr3 that is found only in the Cmr complexes (Fig. S1C). We termed this structure the “stalk loop” (R15–A31). The loop, together with residues in the ZnF domain of Cmr2, engaged in multiple interactions with the bases (Fig. 1G, Fig. S1C). G-5 and A-4 were engaged in multiple interactions with Cmr2R459, which is strictly conserved (Fig. S1C), and Cmr3E19 and Cmr3E21. Further, Cmr3E21, Cmr3F20, and Cmr3P22 directly interacted with the bases of A-4 to A-2. In addition, Cmr3L23 and Cmr3I24 interacted with a group of amino acids building the ZnF domain (Cmr2R218, Cmr2R510, Cmr2G508, Cmr2P507, and Cmr2T465), while Cmr3G26, Cmr3S27, Cmr3H28, and Cmr3A30–31 residues associated with E616 and residues 866–872 of the Cmr2 D2 and Palm2 domains, respectively. Consequently, the stalk loop of Cmr3 was located at a crossroad linking the ZnF, D2, Palm1, and Palm2 domains of Cmr2 with the 5′-RHS of crRNA (Fig. 1E,1G).

On the opposite side of the complex, the crRNA 3′-end was capped by Cmr6 and Cmr1. SisCmr1 is much larger than its orthologs and is only present in Cmr complexes (Fig. 1B-C), and consists of three domains that formed a basic cavity crossed by crRNA, where the protein-RNA contacts involving the phosphate backbone are predominant (Fig. S5E). In addition, two Cmr7 dimers protected the 3′-end of crRNA, this subunit displays a dimeric structure with only 2 homologues (PDB:2X5Q, PDB:2XVO), both found in *S. solfataricus*, which also form dimers (Fig. S6, see below).

### The autoimmunity protection mechanism is revealed by the NTR-bound structure

To understand how SisCmr-β distinguishes between own RNA transcripts and those from invader elements, we determined the cryoEM structure of the NTR-bound SisCmr-β at 3.1 Å global resolution (Fig. S3-S4, STAR Methods Table S2) (Fig. 2A-D). To avoid dissociation of the cleaved target RNA, we used Cmr4 subunits with a D31A substitution to inactivate cleavage (Rouillon et al., 2018; Wang et al., 2019). The structure shows a well-ordered RNA duplex (Fig. 2C-D), and as anticipated the 5’-RHS and 3’-PFS were hybridized. The β-thumb of the Cmr4 subunits was inserted into the RNA duplex, hindering base paring every 6-nt. The assembly of the crRNA-target RNA duplex dissociated the interactions of the catalytic D31 residues in the Cmr4.4 and Cmr4.3 subunits, thus modifying the conformations of the loops where they reside and reorganizing the network of interactions of Cmr4.4D31 with the Palm2 and D4 domains, and those of Cmr4.3D31 with the D4 domain and the Cmr5.3 subunit (Fig. 2D, Fig. S5B). In contrast, upon target RNA binding, the active site loop conformations of Cmr4.1 and Cmr4.2 were also modified but maintained similar interactions with the Cmr5.1 and Cmr5.2 subunits (Fig. 2E, Fig. S5C-D). The Cmr4.2D31 carboxyl was positioned approximately 3.9 Å from the 5’-O of the scissile phosphate, and the surrounding NTR nucleotides were kept in place primarily by salt bridges to residues K145 and R35 in Cmr5.2, and also to Cmr5.3 K56 and base stacking with Cmr4.2 W227 (Fig. 2E). A major rearrangement of Cmr2 was observed between the apo and the NTR structures (Fig. 2F). The Cmr2 protein rotated around the axis formed by the helical filament by ~7° and was displaced by ~6.5 Å.

**Figure 2.**
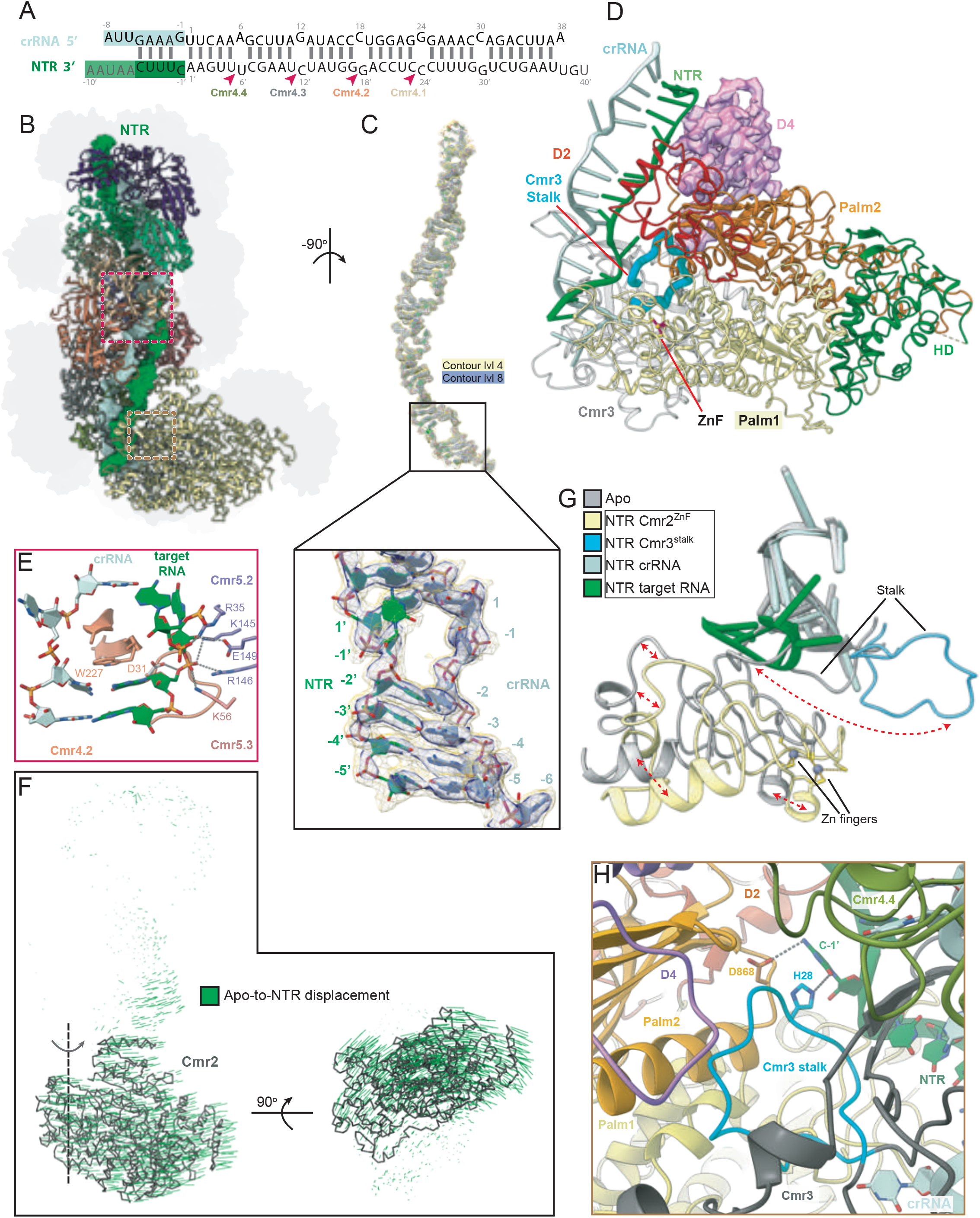
The autoimmunity protection mechanism is revealed by the NTR-bound structure. **(A)** Schematic representation of the crRNA-NTR duplex. A prime (’) is used to distinguish the NTR bases from the crRNA bases. The base pairs observed in the NTR-bound complex are depicted by lines; the observed bases in black, and the bases of the NTR strand that were not traceable are depicted in grey and light shade of green. The crRNA 5’-tag and NTR 3’-anti-tag are shown in teal and green, respectively. Cleavage positions of the target RNA are indicated by red arrow-heads, and the respective Cmr4 subunits providing the active site for the cleavage are indicated (color coded as in fig. 1C). **(B)** The overall structure of the NTR-bound complex with the Cmr7 subunits shown in a flat grey representation; the core subunits in cartoon representation; and the RNA duplex in surface representation (colors are as in Fig. 1C). Red and brown dashed boxes relate to focused views in fig. 2E and 2H, respectively. **(C)** The structure and density of the NTR-crRNA duplex. Close-up view of the base paring between the NTR nucleotides position 1’ to −5’ and crRNA nucleotides 1 to −6. NTR anti-tag nucleotides beyond nucleotide –5′ were not visible in the map. **(D)** Upon NTR binding, and the formation of a 3′-PFS and 5′-crRNA handle duplex, the Cmr3 stalk loop (cyan) undergoes a large rotation into a retracted conformation buried inside a crevice formed between Cmr3 and Cmr2. **(E)** The β-thumb of Cmr4.2 is inserted into the duplex, hindering base paring between crRNA nucleotide 18 and NTR nucleotide 18′. The active-site D31 would be positioned ~3.5 Å from the scissile phosphate, measured from the nearest oxygen to the 5′O of the phosphorous atom. The surrounding NTR nucleotides are kept in place primarily via salt bridges with lysines and arginines of Cmr5.2, but also Cmr5.3 K56, and base stacking with Cmr4.2 W227. **(F)** The NTR-bound structure was superimposed on the apo structure by aligning the Cmr4 subunits. Vectors (green) showing movements of the core subunits relative to the Apo structure were generated using the modevector.py script in PyMOL. The Cmr2 subunit is shown in grey ribbon, and the rest of the structure was omitted for clarity. Vector length correlates with the degree of movement. **(G)** Movement of the Cmr3 stalk (cyan) and Cmr2 Zn finger module (yellow) upon NTR binding (apo form in grey). **(H)** The Cmr3 stalk is positioned at a crossroads, interacting with the Cmr2 Palm1, Palm2, and D4 domains, as well as the cytosine at target RNA position −1′. See also Fig. S1 to S5.

In the Csm complexes, Csm1 is responsible for sensing self *vs*. foreign target RNA through a linker (residues 393–417) and loop L1 (residues 257–265) (You et al., 2019). However, as these regions are not conserved in Cmr2 (Fig. S1B), the self-recognition mechanism in Cmr-β must be different. Accordingly, detailed analysis suggested that the mechanism of self-recognition involves the unique stalk loop in the Cmr3 subunit (Fig. 1E, Fig. S1C). A comparison of the apo and NTR structures revealed that the presence of target RNA induced a large conformational change in the stalk loop, which rotated 90° inwards, and is stabilized by the crRNA–target RNA Watson–Crick base pairing between the 5′-RHS and 3′-PFS (Fig. 2G, Movie S1-S2). This conformational change of the stalk loop is further stabilized by multiple contacts with Palm2 (Cmr2R909/Cmr3G26, Cmr2D876/Cmr3T25, and Cmr2N879/Cmr3E21), ZnF (Cmr2G468/Cmr3E17), and D4 (Cmr2Y1013/Cmr3I24), and C’-1 of the target RNA with Cmr3H28 (Fig. 2H). The movement of the stalk loop unprotected the –5 to –2 region of crRNA and also an area of the ZnF domain. In addition, the Zn^2+^ atom moved 4.7 Å from its position in the apo structure (Fig. 2G). Further, assembly of the RNA duplex was accompanied by the interaction of the target RNA strand with Palm1, and the rigid body movement of the D2 and D4 domains, which packed against the Palm2 domain to permit the entry of the target RNA, thus disengaging the interactions of the catalytic D31 residues in the Cmr4.4 and Cmr4.3 subunits (Movie S3, Fig. S5B-C). All these conformational changes between the apo and NTR complexes were transmitted to the HD domain, leading to the reshaping of its interactions with the Palm1 and Palm2 domains. Although the target RNA was processed at the Cmr4 catalytic centers, no cleavage of ssDNA, UA ssRNA, or production of cOA was detected (Fig. S2E–H), indicating that additional conformational changes were needed to trigger Cmr2 catalytic activities (Wang et al., 2019). Hence, the Cmr3 stalk loop plays a key role in avoiding autoimmunity by Cmr complexes.

### The immunity response mechanism is revealed by a series of CTR-bound structures

To investigate how ssDNA/UA ssRNA degradation and cOA synthesis activities are triggered in the absence of coupling between the 5’RHS and the 3’PFS (Fig. 3A), we determined the structure of the CTR SisCmr-β complex at 3.5 Å global resolution (Fig. S3-S4, STAR Methods Table S2). As in the NTR structure we used the D31A Cmr4 mutant to avoid cleaved target RNA dissociation. We also determined CTR-bound structures in the presence of AMP-PnP to visualize the substrates in the cOA active site. In this sample two discreet states, CTR SisCmr-β/AMP-PnP state 1 (AMPPnP1) and CTR SisCmr-β/AMP-PnP state 2 (AMPPnP2), consistently appeared from focused classification at 2.41 Å and 2.68 Å global resolution. The conformation of the AMPPnP1 structure is very similar to the CTR structure, but with higher resolution information in the AMP-PnP bound sample (Fig. 3B-C, STAR Methods Table S2, Fig. S3-S4). The main differences between AMPPnP1 and AMPPnP2 concerned the Cmr3, Cmr4.4, and Cmr2 area around the cognate target RNA. Although nucleotides –5 to –1 of 3′-PFS were not hybridized with the 5′-RHS, they could be visualized in AMPPnP1 (Movie S4). By contrast, none of them were visible in AMPPnP2 (Movie S5), indicating high flexibility in this region (Fig. 3B-C). The target RNA was observed at the 3′-end, from the +2 nucleotide position onwards, thus looking like the naTR substrate (Fig. 3A–C). In addition, nucleotides U’+6 and C’+7 of the target RNA were not visible because of the movement of the D4 domain, which distorted the duplex and therefore displaced the target RNA in this area (Fig. 3D). This conformational change positioned the conserved Y960 of Cmr2 within a hydrogen bonding distance of Cmr4.4D31, as observed in the apo structure (3.5 Å), suggesting that it would associate with the catalytic D31 residue, disrupting Y960 stacking and stabilizing U’+6 in AMPPnP1 (Fig. 3E). Remarkably, when the naTR (target RNA lacking a 3′-PFS) was used in the activity assays, RNA cleavage was observed, but none of the secondary activities of Cmr2 were detected (Fig. S2E–H), in agreement with the inactive configurations of these subunits in the apo and AMPPnP2 conformation.

**Figure 3.**
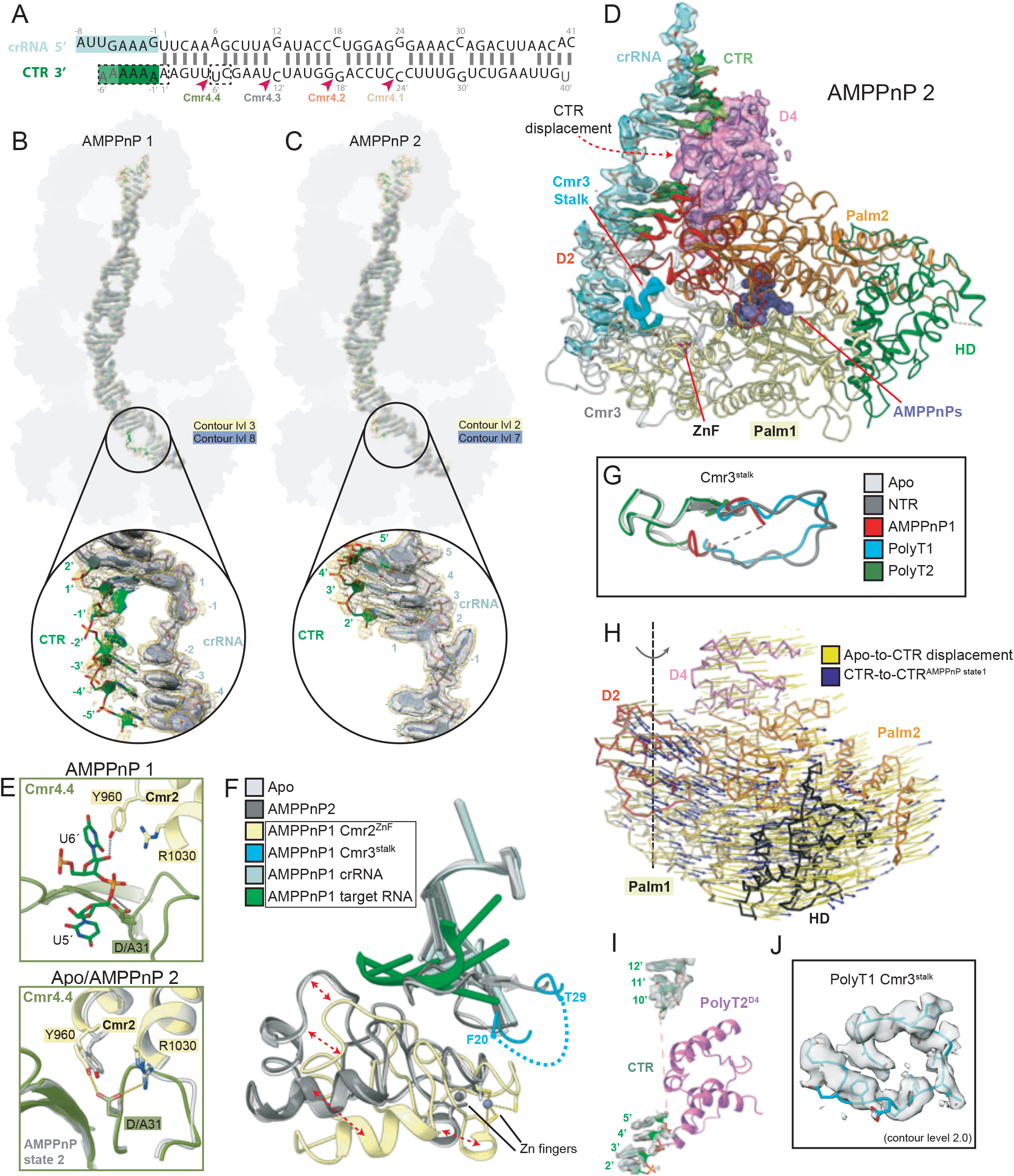
The Immunity Response Mechanism is Revealed by a Series of CTR-Bound Structures. **(A)** Schematic representation of the crRNA-CTR duplex in the CTR-AMPPnP bound state 1 and 2 structures (AMPPnP1 and 2). Observed base pairs are depicted by lines. The crRNA 5’-tag and CTR 3’-PFS are shown in teal and green, respectively. Cleavage positions of the target RNA are indicated by red arrow-heads, and the respective Cmr4 subunits providing the active site for the cleavage are indicated (color coded as in Fig. 1C). Note that four of the six adenine bases of the CTR 3’-PFS were traceable. Bases that could not be traced are highlighted in light shade of green, and dashed boxes, for the AMPPnP1 and 2 structures, respectively. **(B)** and **(C)** The structure and density of the crRNA-CTR duplexes of the AMPPnP1 and AMPPnP2 complexes, respectively. The bound protein is depicted in a flat grey surface representation (Cmr1-Cmr6 in light grey; Cmr7 dimers in dark grey). A close-up view shows the CTR 3’-PFS of the AMPPnP1 structure **(B)** following the same path as the NTR 3’-PFS without forming proper Watson-Crick base-pairing with the crRNA-tag. A close-up view of the AMPPnP2 structure **(C),** with the CTR 3’-PFS not visible and a weak density of bases 2’-5’ (gone at contour level 7). **(D)** Displacement of CTR by the Cmr2 D4 domain as observed in the AMPPnP2 structure. Bases in positions 6’ and 7’ were untraceable, whereas bases 2’-5’ and 8’-12’ could be traced although the density was weak. The Cmr3 stalk loop (cyan) adopts the apo conformation, resting upon the Zn finger module and pressing against the crRNA 5’RHS positions −2 to −5. The densities of the crRNA/CTR (contour level 7) and the D4 domain (contour level 4) are shown in transparent surface representation. Cmr3 is bound at the back of Cmr2. Two molecules of AMPPnP are bound and depicted as purple spheres to show the localization of the cOA site relative to the stalk loop and displaced the CTR. **(E)** Interactions of D/A31 of Cmr4.4, and Y960 and R1030 of Cmr2 in the presence of CTR (top, AMPPnP1 structure) or absence of CTR (bottom, the apo and AMPPnP2 structures superimposed). **(F)** Movement of the Cmr3 stalk and Cmr2 Zn finger module in the AMP-PnP state 1 and 2 structures upon binding of CTR. The AMP-PnP state 2 structure does not exhibit any significant displacement in this region relative to the apo structure, whereas the AMP-PnP state 1 exhibits a displacement similar to that in the NTR-bound structure except for the Cmr3 stalk loop, which could not be traced between F20 and T29. **(G)** Spectrum of the different states of the Cmr3 stalk loop that were observed in the generated structures. **(H)** The CTR and AMPPnP1 structures were superimposed on the Apo structure by aligning the Cmr4 subunits. Yellow vectors indicate the displacement of Cmr2 upon CTR binding (~9° rotation and ~6.5 Å displacement between thecenters of mass), and blue vectors with arrowheads indicate an additional movement of Cmr2 upon binding to AMPPnP (additional ~3° rotation and ~3 Å displacement). **(I)** Structure and density of the target RNA in the vicinity of the Cmr2 D4 domain (purple cartoon) in the PolyT2 structure. Bases in positions 6’-9’ could not be traced. **(J)** The structure and density of the Cmr3 stalk loop of the PolyT1 structure, which is fully traced and has a conformation similar to that observed in the NTR structure.). See also Fig. S1 to S5.

The substantial conformational changes between the apo, NTR, and CTR structures were accompanied by major differences in the Cmr3 stalk loop configuration (Fig. 2G, 3F, Movie S1-S2). The stalk loop was not observed in the CTR-bound or AMPPnP1 structures, indicating lack of the associations observed in the NTR (retracted conformation) and apo (stretched conformation) structures. However, the stalk loop could be traced in a conformation similar to the apo in the AMPPnP2 structure (Fig. 2G, 3F, 3G), suggesting that the alternate conformations of this singular loop of Cmr3 play an important regulatory role controlling the secondary activities of the complex. In addition to the stalk loop and non-complementation in the crRNA handle region, in both the CTR and AMPPnP1 structures, Cmr2 was rotated and translated around the filament axis with respect to the apo (Fig. 3H), as in the NTR-bound complex. However, the extent of Cmr2 rotation around the helical filament axis was ~9° and the displacement was ~6.5Å for the CTR (Fig. 3H), suggesting that the lack of Watson–Crick base paring between the 5′-RHS and 3′-PFS promoted an incremental step of ~2° in the CTR-bound with respect to those observed for the NTR-bound complex. Furthermore, we observed an additional incremental shift of ~2.5° and ~3 Å of the Cmr2 subunit in the AMPPnP1 structure.

To complete the CTR conformational landscape, we analyzed the structure of CTR-bound complex in the presence of AMP-PnP and a 20-nt polyT ssDNA. The sample also displayed the presence of two distinct states after focused classification at 3.0 and 3.1 Å global resolution, which we termed polyT1 and polyT2 respectively (Fig. S3-S4, STAR Methods Table S2). Overall, the polyT1 and polyT2 configurations resembled the AMPPnP1 and 2 structures. Unfortunately, the ssDNA was not observed in the HD catalytic site, indicating that it was largely flexible or unbound. Nevertheless, the structures revealed some important features of the 5′-RHS region where the stalk loop was located. In polyT2, the stalk loop maintains a similar conformation as in the apo and AMPPNP2 structures, building a β-sheet with the conserved β-thumb of Cmr3 (Fig. 1G, 3F-G). The 3′-PFS nucleotides were not visible, and the D4 domain of Cmr2 was packed against Cmr4.4 displacing the nucleotides U’+6 to G’+9 of from the RNA duplex (Fig. 3I), thus resembling the AMPPnP2 structure (Fig. 3D).

In polyT1, the RNA duplex was visualized until the uncoupled 5′-RHS and 3′-PFS region, as in the AMPPnP1 state, and the stalk loop adopted a conformation similar to that in the NTR structure (Fig. 3G, 3J). However, the stalk loop was not stabilized by the interaction of Cmr3H28 with the target RNA, while Cmr2D868 in the Palm2 domain was not associated with the target RNA (Fig. 2H). Collectively, these observations of the CTR-bound state suggest that the stalk loop of Cmr3 alternates between an extended and retracted conformation, rotating ~90°. The different structures appear to capture the target RNA and the stalk loop in different configurations, depending on the presence (NTR) or absence (CTR) of Watson–Crick base pairing between the crRNA 5′-RHS and target RNA 3′-PFS regions (Fig. 3K).

### Target RNA catalysis at Cmr4 active sites

Analysis of the structures representing the CTR-bound stage revealed that the active-site loop (residues G21–P33) of Cmr4.1 and Cmr4.2 superimposed near-perfectly in both apo and target RNA-bound structures (Fig. S5C), whereas this loop exhibited a range of different conformations in the Cmr4.3 and Cmr4.4 subunits of the various solved structures (Fig. S5D). The catalytic D31 residues were positioned 3.8–4.9 Å away from the 5′O of the scissile phosphate (Fig. S7). Among the four catalytic D31 residues, the one in Cmr4.1 was the closest and the one in Cmr4.4 was the furthest away from the corresponding 5′O. The catalytic D31 in the Cmr4.1-4.3 subunits were located in a position where they could act as a catalytic acid to donate a proton to the 5′O and form a leaving OH group, as previously suggested (Osawa et al., 2015). To our knowledge, no other residues have been shown to be part of the active site (Guo et al., 2019; Jia et al., 2019c; Osawa et al., 2015; You et al., 2019). However, we observed that E39 and E149 of the Cmr5 subunits were consistently close to the 2′OH of the ribose adjacent to the scissile phosphate in all three active sites formed between Cmr4 and Cmr5 subunits (3.0–4.7 Å) (Fig. S7). This suggested that they might act as a catalytic bases by deprotonating the 2′OH, with an ensuing nucleophilic attack on the adjacent phosphorous followed by a transition state. Interestingly, E40 of Cmr5 in the chimeric Cmr structure is placed in the same position as SisCmr5 E149 (Osawa et al., 2015), and this residue is conserved in SisCmr5 subunit of Cmr-α and *S. thermophilus* Csm2, further supporting the role in the target RNA cleavage mechanism. By contrast, catalysis should proceed differently in the Cmr4.4 site, in agreement with its different configuration and lower efficiency (Fig. S2E, S5B-D).

### The Cmr2 cOA catalytic site

A model for cOA production by *T. onnurineus* (Ton) Csm1 was proposed by a recent study of the Csm complexes (Jia et al., 2019a). Although Csm1 is a Type III-A ortholog of Cmr2, these subunits exhibit substantial divergence on the sequence and structural levels (Fig. 4A, Fig. S1B). In SisCmr2, two molecules of AMP-PnP were bound in a cleft formed by the Palm1 and Palm2 domains (Fig. 4B). The comparison of SisCmr2 and TonCsm1 cOA sites revealed that the acceptor AMP-PnP molecule superposed well, while the position and conformation of the base moiety of the donor AMP-PnP were different (Fig. 4C). In addition, one of the Mn^2+^ cations of SisCmr2 was bound at a different location, coordinated by E882, and several side chains were either not conserved or exhibited a different conformation at each site, compared with the TonCsm1 structure. Meanwhile, the R389 and R311 associated with the acceptor and donor ATP molecules, respectively, were not conserved in TonCsm1 (Fig. 4C).

**Figure 4.**
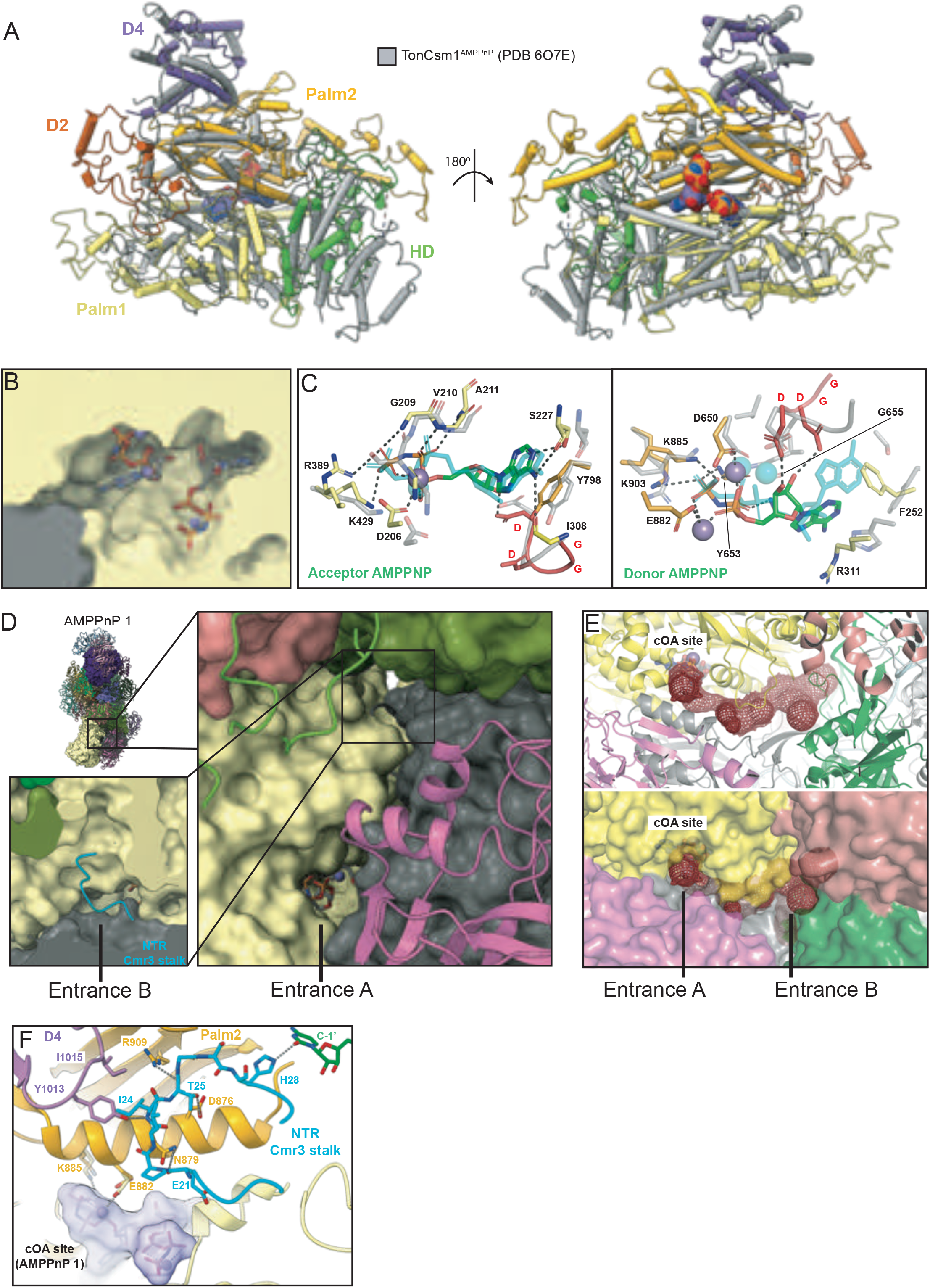
The Cmr2 cOA Catalytic Site. **(A)** Superimposition of SisCmr2 from the AMPPnP1 structure, and TonCsm1-AMPPnP (PDB 6O7E), calculated using the DALI server: RMSD of 4.0, 520 residues aligned, %id 16. **(B)** The ATP-binding pocket of the Cmr2 cOA site, displayed in yellow surface representation with two AMP-PnP moieties and three manganese ions bound (sticks and spheres, respectively). **(C)** The acceptor and donor AMP-PnP moieties and interacting active site residues, left and right respectively, superimposed on the TonCsm1-AMP-PnP. The acceptor AMP-PnP superimposes well, whereas the donor AMP-PnP shows a substantial displacement. **(D)** The two proposed entrances to the ATP binding pocket, A and B, as observed in the AMPPnP1 structure. The Cmr3 stalk loop of the NTR-bound structure blocks entrance B. **(E)** Cavity connecting the entrances A and B with the cOA site (ATP-binding pocket), displayed as a red mesh. The protein is shown in cartoon representation (top view), and in transparent surface representation (bottom view). **(F)** Interaction of the Cmr3 stalk loop with the helix spanning L870-I887 of Cmr2 in the NTR-bound structure, fixates E882 and K885 involved in ATP interactions within the cOA site. The AMP-PnP moieties of the AMPPnP1 structures were superimposed to visualize the localization of the cOA site relative to the NTR Cmr3 stalk loop and relay helix L870-I887. See also Fig. S1 to Fig. S5.

A comparison of the apo, NTR-bound, and different CTR-bound structures revealed two possible entrances to the cOA catalytic center inside Cmr2 (Fig. 4D, E). One of these entrances, entrance A, located at the interface between the α-helix composed by residues E112–N126 of Cmr3 and the Palm2 domain of Cmr2 was clearly observed in the polyT1 and AMPPnP1 structures of the SisCmr-β complex (Fig. 4E). The other accessible path to the cOA catalytic site, entrance B, was located between Palm1, Palm2, a loop of the D4 domain, and the β-sheet formed by the thumb and stalk loop of Cmr3 (Fig. 4E). This cavity is conserved in Csm1 and it has been proposed as the only entry and exit site of the substrate and products in the TonCsm complex (Jia et al., 2019a). We observed that while entrance A was almost entirely occluded in the apo, polyT2 and AMPPnP2 structures, entrance B was occluded by the stalk loop in the NTR complex and polyT1 structures (Fig. 4F). The SisCmr2 helix spanning L870–K885 contributed E882 and K885 to the cOA active site. This helix was kept in place by several interactions with the Cmr3 stalk loop in the NTR structure, suggesting that the NTR-bound complex could not form cOA because of the restriction of movement of crucial active-site residues (Fig. 4F). Therefore, the stalk loop conformation and accessibility appeared to be correlated with the cOA active-site functionality. In structures with the stalk loop in the stretched conformation, entrance A was occluded or narrow, with a relatively more accessible path from entrance B. Conversely, structures in which the stalk loop was in the retracted conformation, entrance A was narrowed and entrance B was occluded (Fig. 4 E). The cavity was open for both entrances only in structures where the stalk loop was not visualized (CTR and AMPPnP1) (Fig. 4E). Collectively, these observations suggested that the interplay between the 5′-RHS and 3′-PFS not only influences the conformation of the stalk loop of Cmr3, but also controls the entrance and exit of molecules to and from the cOA catalytic site, via the stalk loop conformation.

### The Cmr2 HD catalytic center

The HD catalytic center of Cmr2, where ssDNA and UA ssRNA are degraded, resided in a cavity on the subunit surface. The cavity was surrounded by a large electropositive area, favoring interaction with the phosphate backbone of nucleic acids (Fig. 5A-B). The area around the catalytic center contained the conserved H20, D21, D73, and H180 residues; the region of the HD domain that interacted with the rest of Cmr2 was also very well conserved (Fig. 5A-B, Fig S1B). Further, on top of this cavity, a 20-residue loop (G30–K50) was located, forming a lid over the HD domain. The central region of the loop was slightly disordered in the apo and NTR-bound structures, while these residues displayed higher flexibility and fewer residues were modeled into density in the different CTR-bound structures (Fig. 5C). The structural comparison of the apo, CTR, AMPPnP state 1–2, and polyT state 1–2 structures supports the notion that the HD domain undergoes conformational changes, priming it for catalysis (Movies S4, S5). These conformational changes could be allosterically transmitted from the D2–D4 movement to the HD domain via the Palm1 and Palm2 domains, which together with the lack of duplex formation between 5′-RHS and 3′-PFS would activate phosphodiester hydrolysis (Fig. 5D). Considering the location of residues H20 and D21 inside the cavity, we speculate that a conformational change involving the lid on top of the active site is necessary to provide access to the phosphodiester backbone, as supported by the apparent flexibility and electropositive nature of the lid.

**Figure 5.**
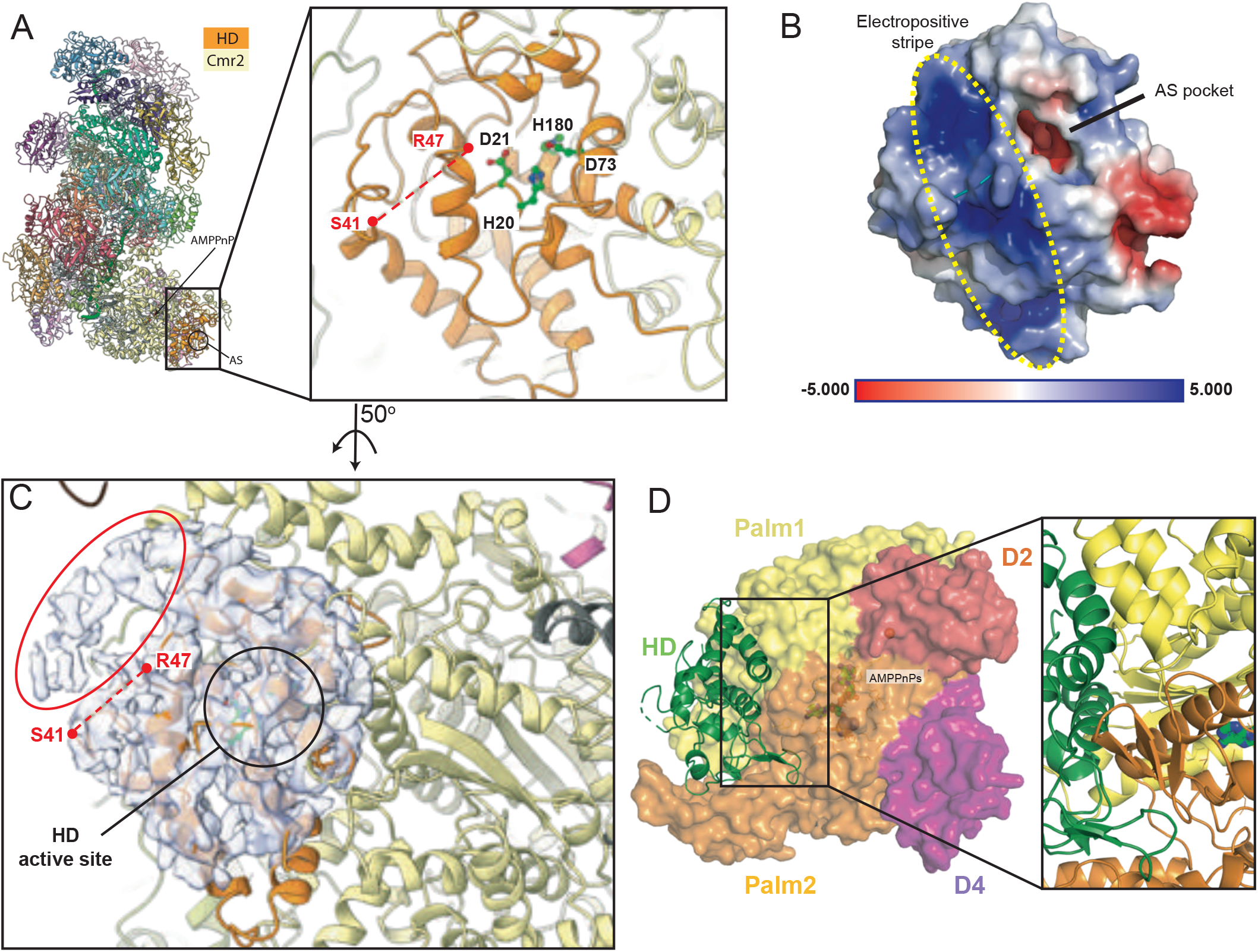
The Cmr2 HD domain. **(A)** Overview of the AMPPnP1 structure, and location of the HD domain and active-site residues in the close-up view. The missing loop just outside the active site pocket (S41 to R47) is indicated with a dashed red line. **(B)** Electrostatic surface potential of the Cmr2 HD domain (calculated using APBS in PyMOL). The active-site pocket and electropositive stripe next to the pocket are highlighted. **(C)** Density map of a portion of the HD domain. A weak unmodeled density highlighted by a red circle, is present next to the missing S41-R47 loop of the HD domain. **(D)** The HD domain, shown in green cartoon representation, is wrapped tightly by the Palm1 and Palm2 domains, shown in yellow and orange representation, respectively. The close-up view shows secondary structural elements of the Palm1 and Palm2 domains involved at the HD interface, through which allosteric changes must be transmitted. The bound AMP-PnP molecules in the cOA site are >15 Å away from the HD domain.

### Allosteric coordination between the Cmr2 active sites

The key catalytic Cmr2 residues are conserved between Csm and Cmr complexes (Fig. S1B), however, divergencies between Type III-A and III-B complexes suggests a different regulatory mechanism. The presence of ATP, the substrate for cOA synthesis, stimulated catalysis in the HD domain of SisCmr-β in contrast to the Csm complex (You et al., 2019). Both ssDNA and UA ssRNA cleavage was enhanced in the presence of the nucleotide, suggesting that the cOA and HD catalytic centers were allosterically connected (Fig S8A-B). A similar activation was observed when AMP-PnP was present; however, AMP-cPP or cOA did not display a stimulation and behaved similarly to the control without activator (Fig S8A). On the contrary, in SisCmr-α, ATP did not enhance cleavage of either substrate (Fig. S8C), suggesting mechanistic differences also between Cmr complexes. Finally, the synthesis of cOA by SisCmr-β only occurs in the presence of Mn^2+^, or in a minor degree when Co^2+^ is present (Fig. S8D).

Our analysis also showed that cleavage of the target RNA is not affected by the abrogation of the catalytic activity in the cOA and ssDNA active sites (Fig S2L). However, the Cmr4D31A mutant displayed a continuous activation of the cOA synthesis and cleavage of ssDNA and UA ssRNA by the HD domain (Fig. S8E-G), suggesting that the inability of the Cmr4 subunits to sever target RNA slowed down the dissociation of the target RNA (Rouillon et al., 2018; Wang et al., 2019), leading to a continuous production of the cyclic compound (Fig. S8E) and cutting of ssDNA and UA ssRNA by the HD domain (Fig. S8F-G). This indicates that the absence of target RNA severance leads to a constant activation of the secondary Cmr2 activities. It also supports the notion that cleavage of the invading target RNA transcript is essential to facilitate its dissociation and switching off the three different activities of Cmr2 (Rouillon et al., 2018).

Amino acid substitution in the HD catalytic center did not abolish the synthesis of cOA. In fact, it promoted increased cOA synthesis (Fig. S2O). However, we observed some differences in the ssDNA and UA ssRNA cleavage when cOA synthesis was abrogated in different complex variants (Fig. S8F-G). The abrogation of cOA synthesis did not abolish ssDNA cleavage, but the severing of ssDNA was affected to a different degree, depending on the cOA catalytic variant tested. Interestingly, stimulation of the ssDNA cleavage appeared to require a hydrolysable phosphodiester between the α and the β phosphate groups of the adenosine triphosphate used as an activator (Fig. S8A). This indicated that the synthesis of cOA allosterically enhances ssDNA cleavage. Finally, the cleavage of UA ssRNA was severely reduced, especially in the Cmr2D650A variant, where the specific cleavage of UA ssRNA was abolished (Fig. S8G), while ssDNA cleavage was reduced (Fig. S8F). Cmr2D650 appeared to be fundamental for cOA synthesis as, based on the structural analysis presented above, it ligated an Mn^2+^ ion coordinating with the donor nucleotide (Fig. 4C), and its substitution hampered catalysis required to generate the cyclic molecule (Fig. S2O. Collectively, the analysis indicated that cOA synthesis by Cmr2 enhanced the catalytic activity in the HD domain.

### Allosteric coordination between Target RNA cleavage and the activation of Cmr2

In the context of the structural information, we explored the coordination between the binding and cleavage of target RNA with the activation of the Cmr2 activities. We designed target RNAs of different lengths, with the 3′-PFS in the CTR ternary complex as the starting point, and tested if their cleavage was linked to the activation of the Cmr2 activities (Fig. 6A–C). One subset of target RNAs contained 6-nt 3′-PFS and incremental segments of the remaining target RNA sequence, while in another subset, only the length of the 3′-PFS was varied (Fig. S8H).

**Figure 6.**
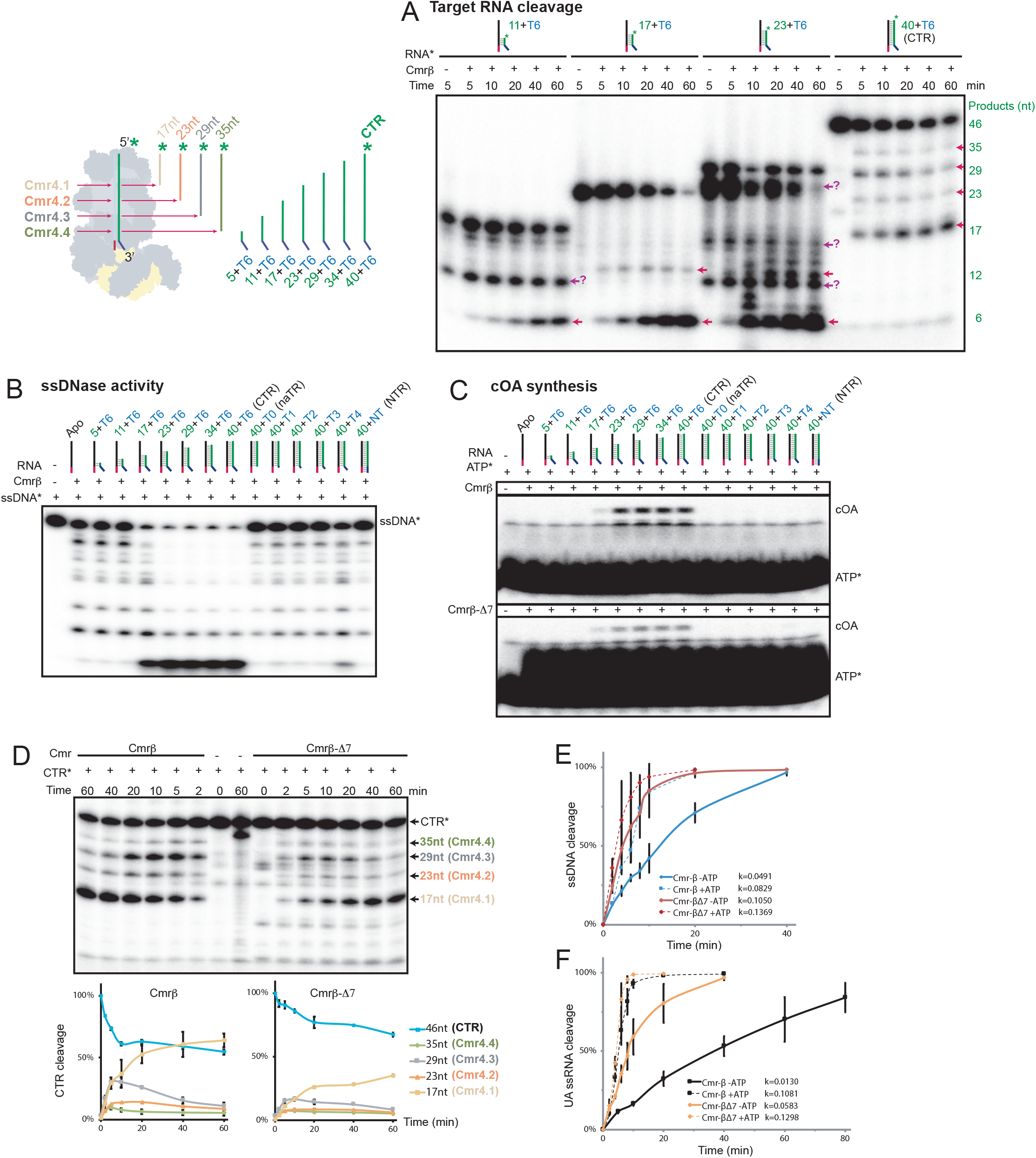
Allosteric Coordination Between Target RNA Cleavage, and the Activation of cOA Synthesis and Nucleic Acid Decay by Cmr2. **(A)** Time-course experiment of target RNA cleavage using RNA molecules of varying lengths. The substrates comprised a 6-mer polyA 3′-PFS (T6) with 11–40 nt of the protospacer region, starting at the +1′ position. The products that were 6-, 12-, and 18-nt in length were expected for the three short target RNAs, whereas those that were 17-, 23-, 29-, and 35-nt in length were expected for the full-length 40+T6 target RNA (CTR). All target RNAs yielded cleavage products of the expected lengths (red arrows). The 11+T6 and 23+T6 target RNAs yielded additional bands that were not accounted for (question marks). Right, a schematic showing the origin of the observed target RNA cleavage fragments, and the short target RNA molecules used in Fig 6A–C on the same scale. The relative position of the Cmr4 subunits in relation to the 3′-PFS and 5′-RHS is also indicated **(B)** ssDNA cleavage by Cmr-β activated by target RNAs of varying lengths, consisting of either a 6-mer polyA 3′-PFS and 5–40 nt of the protospacer region starting at the +1′ position, or a shortened 3′-PFS (T0 to T4) linked to the 40-nt protospacer. ssDNA cleavage is observed in the presence of the 17+T6 target and longer. At least a 4-mer polyA is required to trigger ssDNase activity. **(C)** cOA synthesis by Cmr-β activated by the same set of target RNAs as in Fig. 6B. The cOA synthesis product begins to appear upon the addition of the 17+T6 activator, both in the case of the WT and Δ-7 complexes. **(D)** Target RNA cleavage time-course experiment using the full-length CTR as an activator, comparing the performance of WT and Δ7 complexes. Quantification plots below the gel show progressive CTR degradation and product accumulation. The Cmr4 subunits giving rise to each of the four cleavage fragments are indicated (n = 3 ± SE). **(E)** and **(F)** Quantification plots of ssDNA and UA ssRNA cleavage time course experiments, respectively. Activities are compared for WT and Δ-7 complexes in the presence and absence of ATP (n = 3 ± SE. One of the assay replications related to these data is shown in fig. S8I-J.

The experiment revealed that target RNA was independently cleaved by each of the Cmr4 catalytic centers in the central filament (Fig. 6A). However, ssDNA cleavage was observed only when in addition to the 6-nt of 3′-PFS, the target RNA contained at least 17 additional nucleotides (Fig. 6B, Fig. S8H). Thereby, cleavage of ssDNA appeared to require target RNA binding up to the Cmr4.3 catalytic site, including additional 5′ nucleotides. This suggested that the activation of ssDNA cleavage required not only the lack of complementation between 5′-RHS and 3′-PFS, but also an RNA duplex of a certain length. Such a minimal length of the crRNA-target RNA duplex was necessary for the displacement of the D2 and D4 domains by the coupling of target RNA in that region (Fig. 2D, 3D). Thereby, in the NTR scenario a similar conformational change would prime Cmr2 without leading to its activation (Movie S3), whereas in the CTR scenario, the priming in combination with the uncoupled 3’-PFS would to lead to ssDNA/UA ssRNA cleavage and cOA synthesis (Movies S4-S5).

Since the lack of 3′-PFS (naTR) abrogated all Cmr2 activity (Fig. S2F-H), we asked how many nucleotides of the 3’-PFS were required to achieve Cmr2 activation. Some ssDNA degradation activity was only apparent when 4 of the 6 nucleotides of 3′-PFS were included in the target RNA (40+T4) (Fig. 6B, Fig. S8H). By contrast, in a similar experiment monitoring the synthesis of cOA, we observed that the activity was not fully triggered by the 17+T6 oligonucleotide, and only when 23+T6 target RNA was employed a robust cOA activity was apparent (Fig. 6C, Fig. S8H). Furthermore, a 6-nt 3′-PFS was needed to activate cOA synthesis as none of the target RNAs with a shorter 3′-PFS elicited the synthesis of the cyclic molecule (Fig. 6C, Fig. S8H). These observations suggest that cOA synthesis requires both, the observed D2 and D4 conformational change to prime catalysis, and also a relatively longer 3′-PFS, to allow the uncoupled nucleotides of the 3’-PFS to push the Cmr3 stalk loop away and be sensed by the ZnF in Palm1. Collectively, the data from the above assays fully agree with the progressive rotations and translations observed between the apo, NTR- and CTR-bound structures and the conformational changes observed in the Cmr3 stalk loop and the ZnF area in Cmr2.

Interestingly, we observed that the target RNA cleavage proceeded with different efficiency in the Cmr4 active centers (Fig. 6D), in agreement with the previously observed cleavage (Fig. S2E), and the different active sites configurations in Cmr4.4 and 4.3 compared with those in Cmr4.1 and 4.2 (Fig. S5B-D, S7B). Upon target RNA binding, the interactions of Cmr4.4D31 and Cmr4.3D31 with the Cmr2 and Cmr5 residues, respectively, disappeared. However, Cmr4.4D31 was too far from the 5′O for catalysis (4.9 Å) (Fig. 3E, Fig. S7A-B), in agreement with the low cleavage efficiency observed for this active center (Fig. 6D, Fig. S2E). By contrast, Cmr4.1 and 4.3 were initially the most efficient sites during the cleavage reaction, which correlated well with the notion that both E149 and E39 in Cmr5.1 and 5.3 were located at an appropriate distance so as to facilitate phosphodiester cleavage (Fig. 6D, Fig. S7). Finally, the Cmr4.1 site activity produced the most abundant product band (Fig. 6D), suggesting that the cleaved 5′-end of the target RNA may dissociate faster than the other fragments of target RNA, as also observed for the Csm complexes (Rouillon et al., 2018).

### Cmr7 is a modulator of SisCmr-β activities

We investigated the role of the Cmr7 subunit in the SisCmr-β complex. The Cmr7 dimer contained two centrally positioned cysteines (C63), which are conserved across its known orthologs (Fig. S6I, J). By comparison, Cmr3, Cmr4, and Cmr5 only had one available cysteine each; Cmr6 had two cysteines; and Cmr1 and Cmr2 contained 11 and 13 cysteines, respectively. Interestingly, the Cmr7 dimers exhibited an RMSD of 0.7–1.5 Å and adapted to the different association surfaces along the Cmr-β complex following a general binding pattern, which involved the formation of a specific disulfide bridge between one of the C63 residues of the Cmr7 dimers, and cysteine residues on the surface of other subunits (Fig. S6A). The Cmr7 dimers were distributed around the filamentous core in a regular pattern, following the stacking of Cmr4 and Cmr5 subunits, accordingly (Fig. 1B-C). One Cmr7 dimer was bound to Cmr6, traversing the target RNA-binding groove (Fig. S6E). Two Cmr7 dimers were bound to Cmr2 via the Palm2 domain and wrapping around the HD domain, while one of the Cmr7 dimers was bound to the side of Cmr3 opposite the region of 5′-RHS and 3′-PFS recognition (Fig. S6F). The occupancy of dimers at the 5′-RHS proximal site associated with Cmr3 and Cmr2 was different in the various structures solved. While the apo and NTR structures showed a good occupancy of the Cmr7 dimers in this area, their density was less well defined in the CTR structures. This observation could reflect the notion of a global rearrangement associated with the activation of Cmr2 in the CTR-bound structures (Fig. 1–3) that could affect the association of Cmr7 subunits.

To investigate the role of Cmr7, we generated a SisCmr-β-Δ7 variant which lacked the Cmr7 subunit (Fig S2B), and compared the different activities of WT and Δ7 complexes. SisCmr-β-Δ7 generated twice as much cOA as the WT (Fig. S2I), in agreement with its diminished target RNA degradation activity (Fig. 6D). This deficiency in target RNA cleavage also seemed to promote ssDNA and UA ssRNA cleavage (Fig. 6E, F; Fig S8H, I), in agreement with the behavior of the Cmr4D31A variant deficient in target RNA cleavage (Fig. S2L, S8F-G). Hence, reduced cleavage activity toward the target RNA favored longer residence time of the target RNA in the CTR ternary complex, promoting the other enzymatic activities of Cmr2. Therefore, Cmr7 appeared to allosterically regulate the target RNA cleavage activity of the Cmr4 subunits. However, we cannot discard the possibility that the Cmr7 dimers bound to other SisCmr-β subunits, especially to Cmr2 and Cmr3, could play additional regulatory roles that may be clarified in future studies.

## DISCUSSION

Type III Csm and Cmr complexes display substantial divergences in the effector complex subunit composition, protein orthologues and enzymatic activities. The presence of multiple CRISPR-Cas systems in a genome suggests a combined approach for fighting MGEs. In fact, we speculate that a mechanistic complementation in the immune response between Cmr-α and Cmr-β could happen in *S. islandicus*, as one of them, SisCmr-β, is more active and promiscuous cleaving phosphodiester bonds in both UA ssRNA and ssDNA (Fig. S8B-C), while SisCmr-α generates larger amounts of the cOA secondary messenger (Fig. S2I). Furthermore, we have shown that the Cmr7 subunit modulates the activities of the SisCmr-β complex (Fig. S2I, 6D-F, Fig. S7G-H), and the Cmr7 dimers are located in strategic sites of the complex, i.e associated to Cmr1 at the 3’-end of the crRNA and in the Cmr4.4, Cmr3 and Cmr2 region, where key events of the immune response occur. This led us to speculate whether besides its regulatory role, this subunit could be a structural defense that sterically occludes anti-CRISPR proteins from binding the effector complex (Stanley and Maxwell, 2018). In fact, a new anti-CRISPR protein encoded by the *Sulfolobus* SIRV2 virus against subtype III-B effector complexes has recently been discovered (Bhoobalan-Chitty et al., 2019). However, this anti-CRISPR inhibits the immune response of Cmr-α and Cmr-γ, but it does not target Cmr-β, which is the only Csm/Cmr complex that includes the Cmr7 subunit in its architecture.

### Autoimmunity avoidance

In type III-B Cmr systems, the nascent transcript is captured by the Cmr1 subunit (Li et al., 2017), and then, the effector complex starts to test complementarity of target RNA with crRNA co-transcriptionally at the 3′-end (Amitai and Sorek, 2017; Kazlauskiene et al., 2017). This suggests that testing of coupling between the 5′-RHS and 3′-PFS occurs after Watson–Crick base pairing has produced a long duplex between the target RNA and crRNA and the stalk loop is displaced from the extended configuration (Fig. 1F-G).

In the NTR-bound scenario, the 5′-tag and 3′-antitag hybridize and the assembly of the duplex RNA in this area induces a conformational change of the stalk loop, which retracts and rotates 90° to allow Watson–Crick pairing (Fig. 2G, H). The duplex RNA formed by the 5′-tag and 3′-antitag restrains the possible contacts of the target RNA with the ZnF and Palm1 domains of Cmr2 (Fig. 7A). At the same time, the duplex stabilizes the stalk loop in a retracted conformation that occludes entrance A to the cOA site (Fig. 4D, Fig. 7A), unprotects the ZnF, generating a large movement of this domain, and creates multiple associations of the stalk loop with the Palm2, ZnF, and D4 domains in Cmr2, and with the C’-1 in the 3’-PFS (Fig. 2H, 4F). The assembly of the RNA duplex is also accompanied by the interaction of the target RNA with Palm1, and the rigid body movement of the D2 and D4 domains (Movie S3), which pack against the Palm2 domain to permit the entrance of the target RNA. The changes observed between the apo and NTR structures show that the movement of the stalk loop to interact mainly with Palm2 is also accompanied by a general conformational change of all Cmr2 domains, inducing a rotation and translation of this subunit. Collectively, all conformational changes associated with the entry of target RNA seem to accommodate Cmr2 for cOA synthesis, and ssDNA and UA ssRNA degradation, yet these activities are not triggered because the coupling between the 5′-tag and 3′-antitag favors the retracted conformation of the stalk loop, which in turn arrests Cmr2 activities (Fig S2E–H, Fig. 7A).

**Figure 7.**
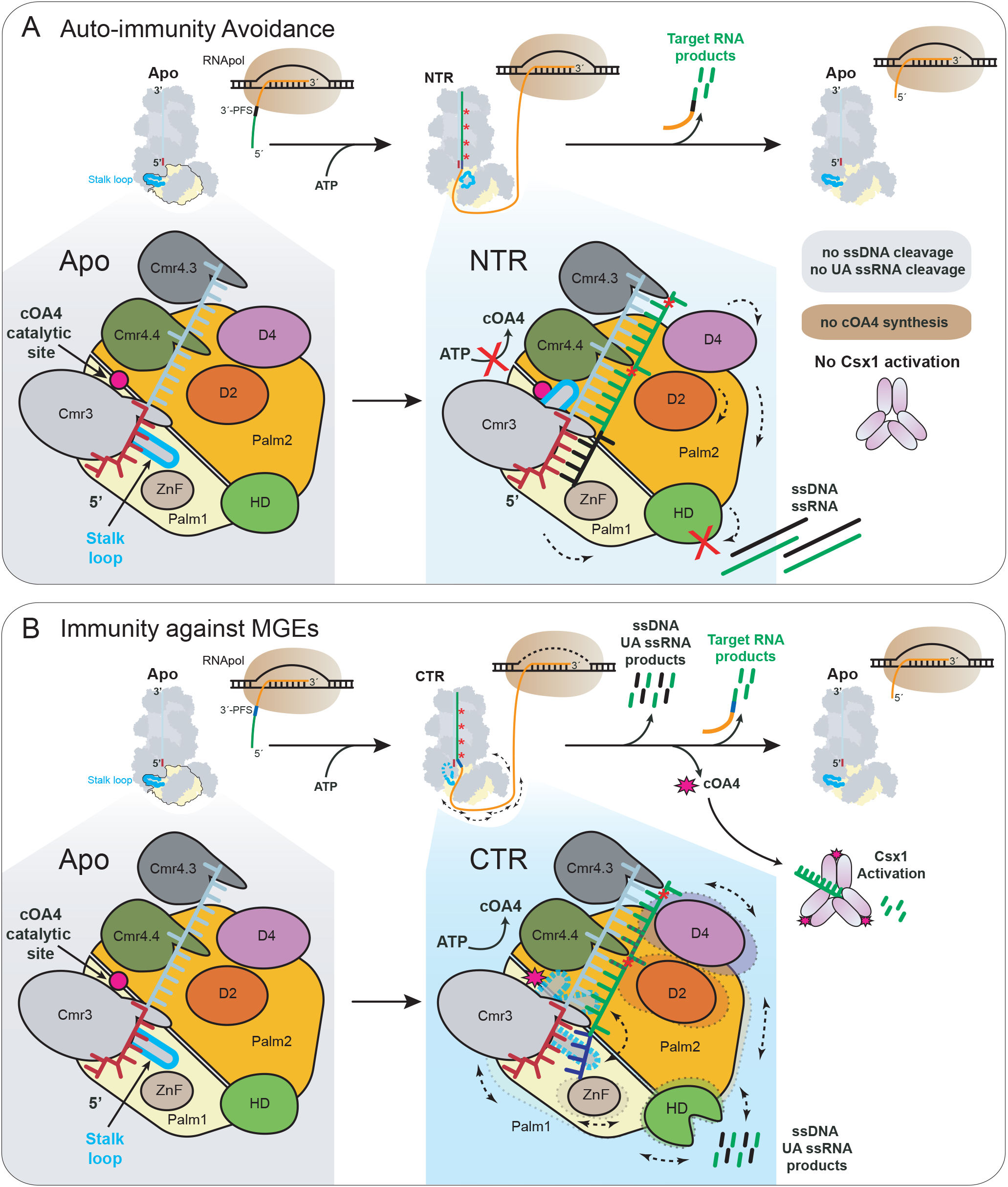
The 3′-PFS Controls the Status of the Cmr3 Stalk Loop, Switching the Immunity On and Off: Model of Type III-B Interference. The crRNA associated with the Cmr complex recognizes a complementary target RNA transcript forming a crRNA-target RNA duplex. **(A)** Auto-immunity avoidance: In case of bidirectional transcription through the CRISPR array, RNA transcripts (green, 3’-PFS in black) complementary to the crRNA spacer (teal) and handle regions (red) may form a crRNA-NTR complex. In this complex, the Cmr3 stalk loop is locked in a retracted conformation (purple), which inhibits ssDNA and UA ssRNA cleavage, and cOA synthesis by the Cmr2 subunit, but target RNA cleavage is promoted, eventually returning the complex to the Apo conformation. **(B)** Immunity against MGEs: In the case of invasion by foreign genetic elements, the RNA is transcribed, and the crRNA will base-pair with the protospacer region (green) but not the 3’-PFS (blue). The Cmr3 stalk loop fluctuates between a range of conformations, allowing ssDNA and UA ssRNA cleavage (in the case of Cmr-β), and cOA synthesis. The auxiliary Csx1 RNase is activated by cOA4, resulting in robust ssRNA interference. Cleavage and dissociation of the target RNA eventually reverts the complex to the apo conformation.

### Activation of the immune response

Comparison of the apo and CTR state structures (CTR, AMPPnP1-2, and polyT1–2) offered molecular snapshots of the different activated conformations in the presence and absence of Cmr2 substrates of the activation state, in which the 5′-RHS and 3′-PFS are uncoupled. These structures displayed a variety of stalk loop conformations, ranging from the extended (apo, AMPPnP2, and polyT2) to the retracted (polyT1), including intermediate states where the loop could not be traced (CTR and AMPPnP1). In addition, the AMPPnP2 and polyT2 structures revealed that the complementation of the crRNA and target RNA is dynamic, especially in the A’+1 to U’+15 region, in which the target RNA needed to displace the D2 and D4 domains of Cmr2, as well as the Cmr5.3 subunit, to allow catalysis in the Cmr4.4 and Cmr4.3 catalytic centers.

A rotation and translation of the Cmr2 structures is observed in the CTR-bound structures where the uncoupled 3’-PFS is visualized (CTR, AMPPNP1 and polyT1), but to a further degree than in the NTR. This movement was observed only in structures in which the loop is flexible or is in the retracted conformation, suggesting that the movement of D2 and D4 induced by the entry of the target RNA primed Cmr2 to a catalytically active state and that the subsequent 5′-tag–3′-antitag Watson–Crick base uncoupling provided an additional step for the initiation of activation (Fig. 7B). This notion was supported by the activity experiments performed with a target RNA lacking the 3′-antitag (naTR), resulting in cleavage of the target RNA but no subsequent ssDNA, UA ssRNA cleavage or cOA synthesis is detected (Fig. S2E–H). The naTR target RNA would correspond to the conformations observed in the AMPPnP2 and polyT2 structures, which did not show the 3′-PFS and displayed the D2 and D4 domains interacting with Cmr4.3 and Cmr 4.4, thus displacing the target RNA in that region. These conformations are similar to the apo, in which catalysis cannot occur. Collectively, the above conformational changes were transmitted toward a compact structure of the cOA catalytic center and a conformational change in the HD domain, which transited from an open to a more compact conformation (AMPPnP1 and polyT1).

### Concluding Remarks

The structural and functional data presented herein indicate that the entry of target RNA induces similar conformational changes in the NTR- and CTR-bound states but without restricting the stalk loop movement in the later scenario. In contrast with Csm complexes where the 3′-PFS is allocated in a binding channel in Csm1 (Jia et al., 2019c; You et al., 2019), we propose an immunity model for the Cmr complexes where the lack of uncoupling with the 5’-RHS facilitates the sensing of the 3’-PFS by the Cmr2 ZnF domain, triggering cOA synthesis and degradation of ssDNA and UA ssRNA (Fig. S1E-H, Fig. 7B). Although Cmr2 is primed in the NTR state, its activities are not unleashed because the stalk loop is stabilized, narrowing entrance A and closing entrance B, and the 3’-PFS does not interact with the ZnF domain (Fig. 7A). Therefore, the stalk loop cannot fluctuate between the stretched and the retracted conformation, which is a requirement to unleash the immune response (Fig. 7B). Finally, cleavage of the target RNA will favor the dissociation of the bound target RNA and new transcript testing.

This work provides new insights into the molecular machinery of SisCmr-β, highlighting the different immune response mechanisms utilized by Type III CRISPR-Cas systems. The large variety of Type III Csm/Cmr complexes and their accessory RNase proteins reflect the diversity and complementation of strategies developed during evolution to adapt the heterogeneous CRISPR-Cas Type III immune systems. Given the large number of prokaryotes containing Type III CRISPR-Cas, new regulatory mechanisms could be identified, and the coordination between different Type III effector complexes simultaneously expressed in the same host organism, could be deciphered in future studies.

## Acknowledgements

We thank the Danish Cryo-EM National Facility in CFIM at the University of Copenhagen for support during cryo-EM data collection. Data processing has been performed at the Computerome, the Danish National Computer for Life Sciences. GM is a member of the Integrative Structural Biology Cluster (ISBUC) at the University of Copenhagen.

## Author Contributions

NS, MF, SS, QS and GM designed the biochemical experiments. YL constructed the mutants, MF, AF optimized the method of Cmr purification, MF, AF, JL, QH purified the wild-type Cmr complexes and the mutated derivatives and conducted biochemical characterizations. MF, AF, JL, QH, QS, SS, NS and GM analyzed the biochemical data. NS prepared the cryoEM samples and EM grids, NS and TP collected the cryoEM images and NS solved and refined the cryoEM structures. NS and GM proceeded with cryoEM structure analysis. QS and GM coordinated the biochemical analysis. The rest of the data were discussed and evaluated with all authors. GM coordinated and supervised the project and wrote the manuscript with NS and input from all the authors.

## SUPPLEMENTARY FIGURES

**Figure S1. Sequence Alignments of the Cmr2 and Cmr3 Subunits, Related to Figure 1-4. (A)** Schematic showing the domain borders and composition of the SisCmr2-β protein. Insertion regions in Cmr2-β compared other Cmr2 sequences are hatched out. **(B)** Sequence alignment of SisCmr2-β against type III-A representatives SisCmr2-α, SsoCmr2, PfuCmr2, and type III-A representatives TonCsm1 and SthCsm1. The Cmr2 domains and secondary structure indicated together above the aligned sequences. Residues involved in the HD active center, coordinating to either the donor or acceptor AMPPnPs (ATPs), ligating the ZnF zinc ion, and residues mutated in the presented work, are all indicated according to the legend in the bottom right corner. **(C)** Sequence alignment of SisCmr3-β against type III-B representatives SisCmr3-α, SsoCmr3, PfuCmr3, and type III-A representatives TonCsm4 and SthCsm4. Stalk loop, 5’-handle, and β-thumb, are indicated above the aligned sequences (colored as indicated in the bottom right corner). The alignments were performed with ClustalO and rendered using ESPript 3.0.

**Figure S2. SisCmr-β Cleaves ssDNA, Generates cOA and Additionally Cuts Non-Complementary ssRNA, Related to Figure 1-4. (A)** Location of the type III-B SisCmr-β gene locus. **(B)** SDS-PAGE gel showing the purified SisCmr-β complex, and a urea gel showing an RNA length marker and the crRNA extracted from the SisCmr-β complex. **(C)** SEC-MALS analysis of the SisCmr-β complex. The change in refractive index (dRI) as a function of protein concentration was used to compute the molar masses. The solid line plotted on the right axis corresponds to the dRI chromatogram from the SEC column. The molar masses across the eluting peak are plotted as open circles (for clarity, only every fifth measurements) on the left axis scale, molar mass. The average molecular weight is displayed on the figure. **(D)** Schematic showing the crRNA hybridized with the different target RNAs used, and the cleavage positions resulting in the experimentally observed fragments. The bottom panel shows the sequence and UA cleavage sites of the S10RNA. **(E)** Target RNA cleavage by Cmr-β in the presence and absence of CTR, naTR, and NTR, respectively. The four 5’-labelled fragments produced are indicated, with the color code used corresponding to the color of the Cmr4 subunit in the schematic (Fig 1C) which cleaves at the associated nucleotide position. **(F)** and **(G)** ssDNA cleavage and cOA synthesis, respectively, by Cmr-β in the presence of CTR, naTR, and NTR. 10 nM Cmr-β was used in the cleavage assay, whereas 20 nM was used in the cOA synthesis assay. **(H)** UA RNA cleavage by Cmr-β in the presence of CTR, naTR, or NTR, and using S10RNA as the labeled substrate. **(I)** cOA synthesis by the wt-β, Δ-7, and wt-α complexes, in the presence of CTR and ATP. The inserts show the quantifications of the bands, comparing wt-β with wt-α, and wt-β with Δ-7, respectively. **(J)** Csx1 band shift upon binding of cOA produced by Cmr-β. **(K)** RNA cleavage by SisCsx1 upon activation by cOA derived from either the Cmr-α-CTR or Cmr-β-CTR complex reaction mixtures where the exact cOA product concentration was unknown, but the volumes added (μl) are indicated on the gel. 25 nM purified material from the SisCmr-α reaction was used as a control. **(L)** Cognate target RNA cleavage by various mutants. The 4-D31A denoted the D31A mutant of Cmr4. The 2-H20A/D21A is the Cmr2 double mutant that renders the HD nuclease inactive. The 2-D802A/D803A has the two aspartates of the GGDD motif in the Cmr2 Palm2 domain substituted to alanine. In the 2-D650A and 2-D652A variants a single conserved aspartate of the Cmr2 Palm2 domain is substituted to alanine. **(M)** ssDNA cleavage of the S10DNA substrate by the WT and 2-H20A/D21A mutant complexes. **(N)** ssRNA cleavage of the S10RNA substrate by the WT and 2-H20A/D21A mutant complexes. **(O)** cOA synthesis in the presence of CTR and ATP, by WT and mutant complexes.

**Figure S3. Single Particle Cryo-EM Analysis of the SisCmr-β Apo and CTR-AMPPnP-Bound Complexes, Related to Figure 1–4. (A)** and **(D)** Representative cryo-EM micrographs of the SisCmr-β apo and CTR-AMPPnP bound complexes, respectively, in vitreous ice on UltrAuFoil 0.6/1.0 grids. **(B)** and **(E)** Reference-free 2D class averages sorted by class distribution (top-left to bottom-right). **(C)** and **(F)** Overview of the cryo-EM data processing workflows for the SisCmr-β apo and CTR-AMPPnP bound complexes, respectively. The processing was generally performed in RELION-3.0. Steps performed in cisTEM are indicated.

**Figure S4. Resolution Assessment and Validation of all Cryo-EM Density Maps, Related to Figure 1-3. (A)** Euler angle distribution plots showing the relative orientation of particles in the final 3D reconstructions, and local resolution maps and central slice through the maps. **(B)** Fourier shell correlation (FSC) curves of the final 3D reconstructions.

**Figure S5. Structural Comparisons of SisCmr4-β and SisCmr1-β, Related to Figure 1–4. (A)** Superposition of SisCmr4-β with TonCsm3 (PDB 6O7E). The active site loop, β-thumb, and a central part of the Cmr4/Csm3 proteins are structurally conserved, but only 57 residues were aligned overall, with an RSMD of 1.09 Å (22% sequence identity). **(B)** The four Cmr4 active sites of the Cmr-β Apo structure, colored as in Fig 1C. The densities of the loop on which the D31 resides (G21-P33), and the Cmr5 R35 and K145, and Cmr2 Y960 and R1030 are shown. **(C)** Superpositions of the Cmr4.1 and 4.2 of the Apo, AMPPnP1, and NTR structures. **(D)** Superpositions of the Cmr4.3 and 4.4 of the Apo, AMPPnP1, and NTR structures. **(E)** Superposition of the Cmr1 subunits of the Apo and AMPPnP1 structures. Despite binding of the CTR, the RMSD is 0.642 covering 462 out of 476 residues.

**Figure S6. Binding of SisCmr7-β Dimers to the Core Complex, Related to Figure 1**. Each Cmr7 dimer and each core subunit contributes one cysteine to a disulfide bridge linking the two units. The Cmr7 dimer has two centrally positioned cysteines. Cmr3, Cmr4, and Cmr5 subunits only have one available cysteine, whereas Cmr6 has 2 cysteines. Cmr1 and Cmr2 have 11 and 13 cysteines, respectively. **(A)** Overview showing the localization of the various Cmr7 dimers decorating the Cmr-β apo core complex (surface and cartoon, respectively). Cmr7 dimers are colored according to the identity of the subunit they bind. **(B)** Four Cmr7 dimers are bound to the core complex via Cmr4 (red surface). The Cmr7.4 dimers stack upon each other in a helical arrangement. The Cmr7 cysteines available for disulfide bridge formation are shown in sphere representation. **(C)** Three Cmr7 dimers are bound to the core complex via Cmr5 (green surface). The Cmr7.5 dimers interact in a different manner compared to Cmr7.4, although still in a regular pattern. **(D)** Two Cmr7 dimers are bound to Cmr1 (blue surface). They do not interact with each other. **(E)** One Cmr7 dimer is bound to Cmr6 sharing the largest interface area of any Cmr7 dimers bound to the core complex (1113 Å2) (yellow surface, cartoon). This dimer traverses the target RNA binding cleft and interacts with Cmr7.5.1, as shown in the inset, and is further stabilized thorugh interactions with Cmr1 (582 Å2). **(F)** Two Cmr7 dimers are bound to Cmr2 (brown surface). Inset shows Cmr7 dimers in cartoon representation in two orientations, binding via the Palm2 domain (yellow surface), and wrapping around the HD domain (green surface). The location of the HD active site is indicated. Although Cmr2 cysteines 721 and 836 are located close to the cysteines of the Cmr7.2.2 and Cmr7.2.1 dimers, respectively, these disulfide bridges could not be modelled convincingly. **(G)** Close-up view of the disulfide bridge, and residues within 5 Å, linking the Cmr7.4.2 dimer and Cmr4, and the density map of this region. **(H)** Superimposition of all Cmr7 dimers from the Apo complex. The dashed circle indicates the position of the Cys63 and Cys63’. **(I)** Superimposition of the two Cmr7 homologues from S. solfataricus (PDB 2X5Q and PDB 2XVO) with Cmr7.4.2. with RMSD of 1.91 Å (green) and 4.17 Å (not shown), respectively. Most secondary structural elements conserved are conserved in the superimposition with of 2X5Q. **(J)** Close up view of the conserved PCD residues (in sticks) located on the center of the concave binding surface. In the bottom left is shown a highly conserved portion of a sequence alignment of Cmr7 from S. islandicus (REY15A) and S. solfataricus (Sso1986 and Sso1725), at the PCD motif (residues 62-68).

**Figure S7. Target RNA Cleavage at the Four Cmr4 Active Centers, Related to Figure 1, 3, and 6. (A)** Overview schematic showing the origin of the 4 target RNA cleavage fragments – 17, 23, 29, 35 nt respectively – observed when providing the complex with a 5’-labelled target RNA. Dashed boxes show the distances of the active site residues in the four Cmr4 active sites with respect to the functional group they target (AMPPnP1 structure), color coded according to the subunit they belong. **(B)** Close-up views of the Cmr4 active sites of the AMPPnP1 structure, color coded according to the schematic in Fig. S7A. **(C)** The Cmr4 active sites of the bottom two subunits (Cmr4.3 and Cmr4.4) in the absence of bound target RNA.

**Figure S8. Deciphering the Role of Cmr7 and Key Residues on Cmr2 activities, Related to Figure 4 and 6. (A)** ATP and AMP-PnP, but not AMP-cPP and cOA4, stimulates cleavage of the single stranded S10DNA, indicating that cOA4 biosynthesis needs to take place in order to allosterically activate the ssDNase activity. **(B)** UA ssRNA (red) and ssDNA (blue) cleavage experiments using the S10RNA and S10DNA substrates respectively, and the CTR-activated Cmr-β in the presence and absence of ATP. The gels shown are single point experiments, whereas the quantification plots are based on time course experiments. N = 3 ± SE. **(C)** CTR-activated Cmr-α does not cleave ssRNA (orange), and ssDNA (green) is cleaved but not stimulated in the presence of ATP. **(D)** cOA synthesis experiment in the presence of different divalent cations. **(E)** cOA synthesis time course experiment in the presence of CTR, ATP, and either the WT or Cmr4 D31A mutant Cmr-β complexes. The insert is a quantification plot based on the gel. **(F)** and **(G)** ssDNA and ssRNA cleavage experiments, respectively, in the presence and absence of ATP, comparing the wt-β complex with the mutants (incubated for 10 minutes). **(H)** The various target RNA sequences, including the short RNAs used in Fig. 6A-C, aligned with the crRNA. The four Cmr4 cleavage sites are indicated on the sequences, and the respective Cmr4 subunits are shown (color coded as in fig. 1B-C). **(I)** and **(J)** ssDNA and ssRNA cleavage time course experiments, respectively, in the presence and absence of ATP, comparing the wt-β-CTR complex (left panels) with the Δ-7 mutant (right panels). The gels are one of 3 replicates used for the quantification plots in fig. 6E, F.

## SUPPLEMENTARY MOVIE LEGENDS

**Movie 1.-** Detailed view of the conformational changes observed in the Cmr3 stalk loop (cyan)

**Movie 2.-** Detailed view of the conformational change of the stalk loop (cyan) in the Cmr3 subunit after the entrance of the target RNA (pink) in the NTR-bound complex (Domains are colored like in fig 1E). Loop the video to better appreciate the conformational change.

**Movie 3.-** Conformational transition between the apo and the NTR-bound complexes in the Cmr3 and Cmr2 area (Domains are colored like in fig 1E). Loop the video to better appreciate the conformational change.

**Movie 4.-** Conformational transition observed between the apo and the CTR-bound AMPPnP1 structures (Domains are colored like in fig 1E). Loop the video to better appreciate the conformational change.

**Movie 5.-** Conformational transition observed between CTR-bound AMPPnP1 and AMPPnP2 structures (Domains are colored like in fig 1E). Loop the video to better appreciate the conformational change.

