## Supplementary material for "Structures of the Cmr-β Complex Reveal the Regulation of the Immunity Mechanism of Type III-B CRISPR-Cas": Methods

**STAR METHODS**

*Strains, growth conditions and transformation of Sulfolobus strains*

*Sulfolobus islandicus* REY15A was isolated from an enrichment culture originated from a hot spring in Iceland (Guo et al., 2011) from which *S. islandicus* E233 and E233S1 were constructed and employed as hosts for genetic manipulation (Peng et al., 2017). *Sulfolobus* strains employed in this work were grown in SCV (0.2% sucrose, 0.2% casa amino acids, 1% vitamin solution plus basic salts) or SCVy (SCV plus 0.05% Yeast extracts) medium at 78° C. If required, uracil was supplemented to 20 μg/ml. Transformation of *Sulfolobus* cells was performed by electroporation as previously described (Deng et al., 2009).

*Construction of plasmids*

To construct a plasmid for expression of the Cmr6β-His recombinant protein for Cmr-β complex co-purification, the *S. islandicus* *cmr6β* gene was cloned into pSeSD1, the *Sulfolobus* expression vector (Peng et al., 2012). However, the *cmr6β* gene carried a NdeI site that had to be removed before insertion of the gene into the NdeI site of pSeSD1, the only usable restriction site for obtaining a C-terminal His-tagged Cmr6β. We employed SOE-PCR (Higuchi et al., 1988) to remove the NdeI site. Two primer sets were designed, including F-NdeI/ R-mut and F-mut/ R-NotI (Table S3), and the designed *cmr6β* gene fragments were obtained by PCR with the above primer sets, using Phanta Max Super Fidelity DNA polymerase (Vazyme) and with the genomic DNA of *S. islandicus* REY15A as the template. A subsequent PCR with the two overlapping fragments as template yielded the designed *cmr6β* gene. The PCR product was then cleaved with NdeI and NotI and the resulting restriction fragment was inserted into pSeSD1 at the NdeI and NotI sites, giving pSe-cmr6β-His. The *cmr-6β*-His fusion gene was then amplified from the plasmid by PCR, using the primer pair of MCS-fwd and MCS-rev (Table S3). The resulting PCR product was digested with SmaI and XhoI and inserted into the SalI and SmaI sites of pAC10-SS1, a CRISPR plasmid that was designed for gene silencing and previously employed for the purification of the *S. islandicus* Cmr-α-RNPs (Han et al., 2017) (Supplementary Table S2), and this yielded pAC-cmr6β. All other pAC-cmrXβ plasmids (pAC-2βD650A and pAC-2βD652A) were constructed with the same strategy using the oligonucleotides listed in Table S3. Several genome-editing plasmids were constructed as described previously (Li et al., 2016) including pGE-4βD31A, pGE-2βHD, pGE-2βpalm2 and pGE-2β (Key Resources Table), and the primers employed for plasmid construction are listed in Table S3. All primers employed in this work were synthesized from Tsingke (Wuhan, China) or TAG Copenhagen A/S (Copenhagen, Denmark). Sequences of all plasmid constructs were verified by DNA sequencing at Tsingke (Wuhan, China) or the MacroGen Europe (Amsterdam, Netherlands).

*Purification of Cmr-β crRNA ribonucleoprotein complex*

The Cmr6-His co-purification procedure developed for *S. solfataricus* and *S. islandicus* Cmr ribonucloprotein (RNP) (Han et al., 2017; Zhang et al., 2016) was adopted for the purification of *S. islandicus* Cmr-β RNPs. *S. islandicus* strains carrying pAC-cmrXβ were grown in SCVy medium at 78° C up to A600 = 0.6-1.0, and cells were collected from at least 6 L of culture for each strain by centrifugation. Cell pellet was re-suspended in Buffer A (20 mM HEPES pH 7.5, 30 mM Imidazole, 500 mM NaCl) and disrupted by French press, followed by centrifuging for 30 min at 12 000 rpm. The supernatant was loaded onto a 1 mL pre-equilibrated HisTrap HP column (GE Healthcare), washed with 30 ml Buffer A, followed by gradient elution into Buffer B (20 mM HEPES pH 7.5, 500 mM Imidazole, 500 mM NaCl). Sample fractions were analyzed by sodium dodecyl sulphate-polyacrylamide gel electrophoresis (SDS-PAGE), and those containing Cmr-β effector complexes were pooled together, concentrated and further purified by size exclusion chromatography in Buffer C (20 mM Tris-HCl pH 8.0, 300 mM NaCl) with a Superdex 200 Increase 10/300 GL column (GE Healthcare). After SDS-PAGE analysis, fractions containing the complete set of Cmr-β subunits were used for further analysis.

*Extraction and analysis of crRNA*

100 μl purified Cmr-β complex was mixed with 200 μl TRIzol (Sigma) and 100 μl chloroform and vortexed to assure disassembly of the complex. Following centrifugation at 12000 g for 5 min, the upper phase was pipetted to a new tube and mixed with 100 μl of chloroform and centrifuged again. The upper phase was then mixed with 1 volume isopropanol and placed at -20° C for 1 hour. All liquid was discarded after a 30 min centrifugation at max speed and the crRNA pellet was washed twice with 70 % cold ethanol. After removing the ethanol, the pellet was air-dried and resuspended in DEPC-treated water. An aliquot of the purified crRNA was 5’-labeled with γ-^32^P-ATP (PerkinElmer) using T4 polynucleotide kinase (New England Biolabs) and separated on a 12% denaturing gel. The labeled RNAs were identified by exposing the gel to a phosphor screen (GE Healthcare) and scanned with a Typhoon FLA 7000 (GE Healthcare).

*Size-exclusion chromatography–multi-angle light scattering (SEC-MALS)*

SEC-MALS experiments were performed using a Dionex (Thermo Scientific) HPLC system connected in-line to a UV detector (Thermo Scientific Dionex Ultimate 3000, MWD-3000), a Wyatt Dawn8+ Heleos 8-angle light-scattering detector and a Wyatt Optilab T-rEX refractive index detector. SEC was performed using a Superdex 200 Increase 10/300 GL column (GE Healthcare) at room temperature in a buffer containing 20 mM Tris pH 8.0, and 500 mM NaCl. For the analysis, 50 μl of Cmr-β complex was injected at 3.3 mg/ml concentration at a flow rate of 0.5 ml/min. ASTRA (version 6.1.17) software was used to collect the data from the UV, refractive index, and light scattering detectors. The weight average molecular masses, M_w_, were determined across the elution profile from static LS measurements using ASTRA software and a Zimm model, which relates the amount of scattered light to the weight average molecular weight of the solute, the concentration of the sample, and the square of the refractive index increment (dn/dc) of the sample.

*DNA and RNA cleavage assays*

RNA and DNA substrates used in the cleavage assays were purified by recovering the corresponding bands from a denaturing gel and 5’-labeled with γ-^32^P-ATP (PerkinElmer) using T4 polynucleotide kinase (New England Biolabs). ssRNA and ssDNA cleavage assays were conducted in 10 μl reaction mixtures containing 20 mM MES (pH 6.0), 5 mM MnCl_2_, and 20 nM Cmr-β wild-type or mutant complex (unless otherwise indicated in the corresponding figure legends). For target RNA cleavage experiments, 20 nM target RNA was used. For the target RNA-activated ssDNA or ssRNA cleavage assays, unlabeled target RNA was supplemented to 200 nM, and 50 nM ssDNA (S10DNA) or RNA (S10RNA) substrate was used. For the ATP stimulated cleavage assays, 1nM α^32^P-ATP and 100nM cold ATP was added to the reactions. All cleavage reactions were performed at 70° C for the 20 minutes unless otherwise indicated in the figure legends, and the reaction was stopped by the addition of 2x RNA loading dye (New England Biolabs) and kept on ice. Immediately before loading, reaction samples were heated for 5 min at 95° C and analyzed by denaturing polyacrylamide gel electrophoresis using a 18% gel. Cleavage products were visualized by phosphor imaging. RNA ladders were generated by Decade™ Marker RNA (Ambion) following the instructions, while 10/60 DNA ladders were purchased from IDT and ^32^P-labeled with T4 polynucleotide kinase.

*Cyclic oligoadenylate (cOA) synthesis assays*

Synthesis of cyclic oligoadenylates (cOAs) by the Cmr-β was performed as previously described (Han et al., 2018). In brief, the reaction containing 20 mM MES pH 6.0, 5 mM MnCl_2_ (unless otherwise indicated), 1 nM α^32^P-ATP (PerkinElmer) and 100 nM cold ATP, 200 nM CTR and the 20 nM Cmr-β, Δ7-β mutant, or Cmr-α was incubated at 70° C for 20 minutes unless otherwise is indicated, and stopped by chilling on ice and adding 2x RNA loading dye (New England Biolabs), followed by 24% denaturing gel analysis.

*In vitro SisCsx1 binding assays*

Preparation of the radioactively labelled cOAs using ^32^P-α-ATP was performed with 0.5 μl of 3000 Ci/mmol ^32^P-α-ATP (PerkinElmer) and 10 μM cold ATP, 200 nM CTR and 1 μM SisCmr-β in a reaction buffer (20 mM MES pH 6.0, 5 mM MnCl_2_) and incubated at 70° C for 40 min. The reaction was separated on a 24% denaturing gel (19:1 acrylamide:bis-acrylamide in TBE with 8 M urea). The cOA_4_ was extracted from the gel by overnight incubation of the gel slice in MilliQ water. The binding assay was performed by adding 2 μl ^32^P-labelled cOA to increasing amount of SisCsx1 (0, 8, 16, 32, 64, 128, 256 nM in 20 mM MES pH 6.0, 5 mM MnCl_2_ and incubating at 70° C for 5 min, followed by analysis on a 10% TBE native gel (Invitrogen), and visualization by phosphor imaging.

*In vitro RNA cleavage assay by SisCsx1*

The RNA cleavage assay was performed with 18 nM SisCsx1, 2.5 μM 5′FAM-labeled RNA2, and increasing amounts of labelled cOA in a reaction buffer (20 mM MES pH 6.0, and 5 mM MnCl_2_). The reactions were incubated at 70° C for 5 min and stopped by adding 2x stop buffer (8 M urea and 100 mM EDTA at pH 8.0) and cooling on ice. Samples were loaded on a 15% Novex TBE-urea poly-acrylamide gel (Invitrogen), and visualized using an Odyssey FC Imaging System (Li-Cor).

*Cryo-EM sample preparation and data collection*

The wild-type SisCmr-β binary (apo) complex was concentrated to ~4.5 mg/mL, diluted in buffer containing 150 mM NaCl, 10 mM Tris-HCl pH 8, and kept on ice for cryo-EM specimen preparation. In order to obtain the higher order ternary complexes of substrate-bound *Sis*Cmr-β, the Cmr4 D31A variant of the *Sis*Cmr-β binary complex was used instead of the wild-type to avoid back-bone cleavage of the target RNA. The CTR, NTR, and CTR + AMPPnP bound complexes were assembled by mixing in buffer containing 250 mM NaCl, 20 mM MES pH 6, 1 mM MnCl_2_, and 1 mM DTT. The CTR + AMPPnP + polyT-bound complex was assembled in 150 mM NaCl, 10 mM MES pH 6, 1 mM CaCl_2_ in order to inhibit ssDNase activity, and DTT was excluded to avoid reduction of disulphide bridges. The mixed samples were subsequently incubated for 20 minutes at 50° C. The final concentrations of samples applied for cryo-EM were 0.8 mg/mL Cmr-β binary complex, 1.4x molar ratio target RNA and polyT, and 0.2 mM AMPPnP.

3 µL sample was applied to UltrAuFoil 300 mesh R0.6/1.0 holey grids (Quantifoil) glow-discharged for 30 s at 5 mA (Leica EM ACE200), and plunge-frozen in liquid ethane, cooled with liquid nitrogen, using a Vitrobot Mark IV robot (FEI, Thermo Fisher Scientific) with the following settings: blotting for 3-4 s, 100% humidity and 4° C. Cryo-EM micrographs were collected on at Titan Krios G3 transmission electron microscope (FEI, Thermo Fisher Scientific) operating at 300 kV at liquid nitrogen temperature. The datasets (dose-fractioned movies) were acquired using a Falcon 3ED Direct Electron Detector (FEI, Thermo Fisher Scientific) in electron counting mode, except for the Cmr-β-NTR ternary complex which was collected in linear mode. A nominal magnification of 96,000 was used resulting in a calibrated pixel size of 0.832 Å. A single exposure was collected per hole, with an applied defocus range of approximately -1.7 to -2.6 µm. For the Cmr-β, Cmr-β-CTR, and Cmr-β-CTR-AMPPnP-polyT specimen, the exposures consisted of 40 frames and a dose rate of 1.0 e^-^/Å^2^. For the Cmr-β-CTR-AMPPnP specimen, the exposures consisted of 50 frames and a dose rate of 0.8 e^-^/Å^-2^, and for the Cmr-β-NTR specimen collected in linear mode, the exposures consisted of 19 frames and a dose rate of 4.5 e^-^/Å^-2^. EPU was used for automated data acquisition (FEI, Thermo Fisher Scientific).

*Image processing*

All datasets were initially processed using *cis*TEM-1.0-beta (Grant et al., 2018), and the datasets of the Cmr-β, Cmr-β-CTR-AMPPnP, and Cmr-β-CTR-AMPPnP-polyT specimen were subsequently re-processed using the RELION 3.0 workflow to push for higher resolution reconstructions (Scheres, 2012; Zivanov et al., 2018). Beam-induced drift correction and contrast transfer function (CTF) estimation were performed using Unblur (Grant and Grigorieff, 2015) and CTFFIND4 (Rohou and Grigorieff, 2015), respectively, as implemented in *cis*TEM. Particles were picked from images with thon rings detected at 4 Å and beyond, using the implemented “ab initio” algorithm based on a simple low-pass filtered disc for searching. The particles were extracted and sorted by two rounds of 2D classification. An initial model was generated using the *cis*TEM *ab initio* reconstruction algorithm followed by 3D auto-refinement. Correct handedness was ensured by rigid body fitting of homology models in UCSF Chimera (Pettersen et al., 2004) (see *Atomic Model Building and Refinement*), and the map was used as a reference for subsequent 3D auto-refinements of other datasets.

Briefly, for the Cmr-β apo dataset 143 K coordinates were auto-picked from 3,785 micrographs (binned to 1 Å/pixel), extracted and pruned to 49 K particles by two rounds of 2D classification. These particles were used for 3D auto-refinement, and the resulting map and parameters were used to initiate manual local 3D classification. One of five classes (32 K particles) was selected for the final 3D refinement, resulting in a map with an overall resolution of 3.15 Å based on the gold-standard Fourier shell correlation (FSC) 0.143 cutoff criterion.

In the Cmr-β-CTR complex, 72 K coordinates were auto-picked from 3,785 micrographs (binned to 1 Å/pixel), extracted and pruned to 54 K by 2D classification, which in turn were used for 3D auto-refinement followed by a manual local 3D classification. One of five classes (30 K particles) were selected for the final 3D refinement, resulting in a map with an overall resolution of 3.4 Å.

For Cmr-β-NTR, 344 K coordinates were auto-picked from 5,190 micrographs (binned to 1Å/pixel), extracted and pruned to 189 K particles by two rounds of 2D classification. 3D auto-refinement was followed by a manual local 3D classification, and one of five classes (112 K particles) was selected for the final 3D refinement, resulting in a map with an overall resolution of 3.07 Å.

For Cmr-β-CTR-AMPPnP, 321 K coordinates were auto-picked from 3,912 micrographs (not binned), extracted and pruned to 232 K particles by two rounds of 2D classification. 3D auto-refinement was followed by two rounds of manual local 3D classification, and one out of the final 3 classes (88 K particles) was selected for the final 3D refinement, resulting in a map with an overall resolution of 2.62 Å.

For Cmr-β-CTR-AMPPnP-polyT, 240 K coordinates were auto-picked from 5,298 micrographs (not binned), extracted and pruned to 101 K particles by two rounds of 2D classification. 3D auto-refinement of these particles produced a map with an overall resolution of 3.15 Å, which in turn was locally classified into 3 classes by using a soft focus mask centered on the region corresponding to Cmr2. This resulted in reconstructions ranging from 3.25 to 3.55 Å resolution that exhibited slight variation in the region of the maps corresponding to Cmr2.

For the RELION workflow, individual frames of the micrographs were aligned and dose-weighted using MotionCor2 (Zheng et al., 2017). CTF parameters were estimated using Gctf (Zhang, 2016), and images were discarded according to the assessed maximum resolution (<4 Å) and accuracy of fitting. For the Cmr-β apo dataset, templates for auto-picking were acquired by providing a 3D reference reconstructed in *cis*TEM, which was internally used to generate 2D images by projection. 614 K coordinates were auto-picked and reduced to 101 K through three rounds of 2D classification and particle sorting. 3D auto-refinement was performed with a reference mask and solvent-flattened FSCs, initially resulting in a 3.35 Å reconstruction. Subsequently, a heterogeneous particles subset (5 K) was removed by 3D classification, and iterative rounds of CTF refinement and Bayesian polishing improved the final reconstruction to 2.99 Å. Particle parameters and images were exported to *cis*TEM in order to perform a focused 3D classification with local searches. First, the reconstruction was re-generated with the resolution now estimated to 2.59 Å (using the soft spherical mask approach implemented in *cis*TEM). Next, a focus mask of 30 Å radius centered on the HD domain of the Cmr2 subunit was used in the 3D classification to generate three classes reconstructed to 3.54 Å, 2.86 Å, and 3.19 Å respectively. Class 2 was further refined to 2.75 Å.

For Cmr-β-CTR-AMPPnP, 785 K coordinates were auto-picked and reduced to 389 K through two rounds of 2D classification and particle sorting. 3D classification was initially used to identify a homogeneous complex of 161 K particles, and a large heterogeneous particles subset which mainly suffered from neighboring particles physically touching, and thus, was omitted from further analysis. 3D auto-refinement was performed on the homogeneous subset, initially generating a 3.04 Å map. Two rounds of CTF refinement and Bayesian polishing with a round of particle sorting in between, improved the final reconstruction to 2.7 Å. A reference mask and solvent-flattened FSCs were used in the final 3D auto-refinement. Ewald sphere correction was attempted without yielding any improvements on map resolution. Particle parameters and images were exported to *cis*TEM in order to perform focused 3D classification. First, the reconstruction was re-calculated to 2.34 Å. As above, focused 3D classification and refinement was performed resulting in three reconstructions at 2.68 Å (termed AMPPnP2), 3.0 Å, and 2.41 Å (termed AMPPnP1).

For Cmr-β-CTR-AMPPnP-polyT, 429 K coordinates were auto-picked and reduced to 239 K through two rounds of 2D classification and particle sorting. The data was 3D classified into five classes of which three (136 K) were merged for 3D auto-refinement, resulting in an initial 3.41 Å resolution map. Two rounds of CTF refinement and bayesian polishing with particle sorting based on the refined model sandwiched in between, improved the final reconstruction to 3.08 Å. A reference mask and solvent-flattened FSCs were used in the 3D auto-refinement. Particle parameters and images were exported to *cis*TEM in order to perform focused 3D classification. First, the reconstruction was recalculated to 2.84 Å. Focused 3D classification and refinement was performed resulting in 3.14 Å (termed polyT2), 3.07 (PolyT1) Å, and 3.37 Å reconstructions. Local resolution analysis was performed with MonoRes (Vilas et al., 2018), and directional resolution anisotropy was analyzed using the 3D-FSC server (Tan et al., 2017).

*Atomic model building and refinement*

Homology models of all proteins generated using Phyre2 (Kelley et al., 2015) and I-TASSER (Yang et al., 2015) were placed in an initial ~4 Å reconstruction of the Cmrβ complex using map segmentation and rigid body fitting in UCSF Chimera. A higher resolution map subsequently allowed *ab initio* model building of the helical core by map-to-model implemented in PHENIX (Adams et al., 2010). This model was connected, extended, and rebuilt *de novo* in COOT, using the initial homology model as a guide (Emsley and Cowtan, 2004). Subsequently, the sharpened 2.4 Å Cmr-β-CTR-AMPPnP1 complex map readily allowed model building of the RNA duplex, and roughly 90% of Cmr2 and 30-70% of the individual Cmr7 dimers. Moreover, sharpened and unsharpened maps of the Cmrβ-NTR complex, as well as the multi-chicken tool in COOT, were used to guide model building towards the highest possible completeness of the models of Cmr2 and the Cmr7 dimers. The Namdinator molecular dynamics flexible fitting server was used to generate models fitting the various maps (Kidmose et al., 2019), before manually checking through the maps and refining in phenix.real_space_refine. All model and map images were created using ChimeraX (Goddard et al., 2018), except for the vector models (Fig. 2B, 3H), cavities figure (Fig. 4E), electrostatic potential map (Fig. 5B), and Cmr4 superpositions (Fig. S5C, D).

**KEY RESOURCES TABLE**

| REAGENT or RESOURCE | SOURCE | IDENTIFIER |
| --- | --- | --- |
| Chemicals, Peptides, and Recombinant Proteins | | |
| Tris | Sigma-Aldrich | Cat# T1503-5KG |
| DTT | Sigma-Aldrich | Cat# D0632-10G |
| HEPES | Sigma-Aldrich | Cat# H3375-250G |
| Imidazole | Millipore | Cat# 1047161000 |
| Urea | Millipore | Cat# 1084871000 |
| MES | Sigma-Aldrich | Cat# M0164 |
| MnCl_2_ | Sigma-Aldrich | Cat# 63535 |
| CaCl_2_ | Sigma-Aldrich | Cat# 223506 |
| AMPPNP | Jena Bioscience | Cat# NU-407-10 |
| AMPPCP | Jena Bioscience | Cat# NU-421-10 |
| ATP | Sigma-Aldrich | Cat# A2383 |
| [α-^32^P] ATP | PerkinElmer | Cat# NEG003H250UC |
| [γ-^32^P] ATP | PerkinElmer | Cat# NEG002A100UC |
| TRIzol | Sigma-Aldrich | Cat# 93289 |
| T4 polynucleotide kinase | New England Biolabs | Cat# M0201L |
| Phanta Max Super Fidelity DNA polymerase | Vazyme | Cat# P505 |
| NdeI, NotI, SmaI, SalI, and XhoI restriction enzymes | Thermo Fisher |  |
| 2x RNA loading dye | NEB | Cat#B0363S |
| Critical Commercial Assays | | |
| HisTrap HP | GE Healthcare | Cat# 17-5247-01 |
| Superdex 200 Increase, 10/300 GL | GE Healthcare | Cat# 28-9909-44 |
| Deposited Data | | |
| Map of Cmr1_1_2_1_3_1_4_4_5_3_6_1_7_26_^crRNA^ binary complex | This paper | EMDB: 10102 |
| Map of Cmr1_1_2_1_3_1_4_4_5_3_6_1_7_22_^crRNA^-cognate target RNA ternary complex | This paper | EMDB: 10209 |
| Map of Cmr1_1_2_1_3_1_4_4_5_3_6_1_7_22_^crRNA^-cognate target RNA ternary complex bound to AMPPNP, state 1 | This paper | EMDB: 10117 |
| Map of Cmr1_1_2_1_3_1_4_4_5_3_6_1_7_22_^crRNA^-cognate target RNA ternary complex bound to AMPPNP, state 2 | This paper | EMDB: 10126 |
| Map of Cmr1_1_2_1_3_1_4_4_5_3_6_1_7_22_^crRNA^-non-cognate target RNA ternary complex | This paper | EMDB: 10119 |
| Map of Cmr1_1_2_1_3_1_4_4_5_3_6_1_7_26_^crRNA^-cognate target RNA ternary complex bound to AMPPNP, state 1, in the presence of polyT (20-mer) | This paper | EMDB: 10197 |
| Map of Cmr1_1_2_1_3_1_4_4_5_3_6_1_7_26_^crRNA^-cognate target RNA ternary complex bound to AMPPNP, state 2, in the presence of polyT (20-mer) | This paper | EMDB: 10196 |
| Model of Cmr1_1_2_1_3_1_4_4_5_3_6_1_7_26_^crRNA^ binary complex | This paper | PDB: 6S6B |
| Model of Cmr1_1_2_1_3_1_4_4_5_3_6_1_7_22_^crRNA^-cognate target RNA ternary complex | This paper | PDB: 6SIC |
| Model of Cmr1_1_2_1_3_1_4_4_5_3_6_1_7_22_^crRNA^-cognate target RNA ternary complex bound to AMPPNP, state 1 | This paper | PDB: 6S8B |
| Model of Cmr1_1_2_1_3_1_4_4_5_3_6_1_7_22_^crRNA^-cognate target RNA ternary complex bound to AMPPNP, state 2 | This paper | PDB: 6S91 |
| Model of Cmr1_1_2_1_3_1_4_4_5_3_6_1_7_22_^crRNA^-non-cognate target RNA ternary complex | This paper | PDB: 6S8E |
| Model of Cmr1_1_2_1_3_1_4_4_5_3_6_1_7_26_^crRNA^-cognate target RNA ternary complex bound to AMPPNP, state 1, in the presence of polyT (20-mer) | This paper | PDB: 6SHB |
| Model of Cmr1_1_2_1_3_1_4_4_5_3_6_1_7_26_^crRNA^-cognate target RNA ternary complex bound to AMPPNP, state 2, in the presence of polyT (20-mer) | This paper | PDB: 6SH8 |
| Experimental Models: Organisms/Strains | | |
| *S. islandicus* E233 | (Peng et al., 2017) | N/A |
| *S. islandicus* E233S1 | (Peng et al., 2017) | N/A |
| *S. islandicus* Δcmr-2β | This paper | N/A |
| *S. islandicus* Δcmr-α | This paper | N/A |
| *S. islandicus* Cmr4βD31A ΔIA | This paper | N/A |
| *S. islandicus* Cmr2βHD ΔIA | This paper | N/A |
| *S. islandicus* Cmr2βpalm2 ΔIA | This paper | N/A |
| *S. islandicus* MF1 | Han et al., 2017 | N/A |
| *Escherichia coli* DH5α | Home-made | N/A |
| Oligonucleotides | | |
| RNA and DNA substrates | This paper | Table S2 |
| DNA primers | This paper | Table S3 |
| Recombinant DNA | | |
| pSeSD1 | (Peng et al., 2012) | N/A |
| pSe-Rp | (Peng et al., 2015) | N/A |
| pMID-Cmrα | (Peng et al., 2013) | N/A |
| pAC-MS1 | Han et al., 2017 | N/A |
| pcmr6β | This paper | N/A |
| pAC-cmr6β | This paper | N/A |
| pAC-2βD650A | This paper | N/A |
| pAC-2βD652A | This paper | N/A |
| pGE-IA | (Han et al., 2017) | N/A |
| pGE-4β D31A | This paper | N/A |
| pGE-2β HD | This paper | N/A |
| pGE-2β palm2 | This paper | N/A |
| pGE-2β | This paper | N/A |
| pGE-7β | This paper | N/A |
| Software and Algorithms | | |
| COOT | (Emsley and Cowtan, 2004) | <https://www2.mrc-lmb.cam.ac.uk/personal/pemsley/coot/> |
| PHENIX | (Adams et al., 2010) | <https://www.phenix-online.org/> |
| Namdinator | (Kidmose et al., 2019) | <https://namdinator.au.dk/> |
| ChimeraX | (Goddard et al., 2018) | <https://www.rbvi.ucsf.edu/chimerax/> |
| UCSF Chimera | (Pettersen et al., 2004) | <https://www.cgl.ucsf.edu/chimera/> |
| PyMOL | The PyMOL Molecular Graphics System, Version 2.0 Schrödinger, LLC. | http://www.pymol.org/2/ |
| RELION | Scheres, 2012 | [https://www3.mrc-lmb.cam.ac.uk/relion//index.php/Main_Page](https://www3.mrc-lmb.cam.ac.uk/relion/index.php/Main_Page) |
| cisTEM | (Grant et al., 2018) | <https://cistem.org/> |
| SCIPION | (de la Rosa-Trevin et al., 2016) | <http://scipion.i2pc.es/> |
| MonoRes | (Vilas et al., 2018) | <http://scipion.cnb.csic.es/m/myresmap> |
| MotionCor2 | (Zheng et al., 2017) | <http://msg.ucsf.edu/em/software/index.html> |
| gCTF | (Zhang, 2016) | <https://www.mrc-lmb.cam.ac.uk/kzhang/> |
| 3D-FSC | (Tan et al., 2017) | <https://3dfsc.salk.edu/> |
| ImageQuant TL | GE Healthcare, version 8.1 | https://www.gelifesciences  .com/ |

**Table S1. Statistics of Cryo-EM data Processing and Refinement. Related to figures S3-S4.**

|  | Cmr1_1_2_1_3_1_4_4_5_3_6_1_7_26_β^crRNA^ | Cmr1_1_2_1_3_1_4_4_5_3_6_1_7_22_β^crRNA^-NTR | Cmr1_1_2_1_3_1_4_4_5_3_6_1_7_22_β^crRNA^-CTR-AMPPNP, state1 | | Cmr1_1_2_1_3_1_4_4_5_3_6_1_7_22_β^crRNA^-CTR-AMPPNP, state2 |
| --- | --- | --- | --- | --- | --- |
|  | EMDB: 10102 | EMDB: 10119 | EMDB: 10117 | | EMDB: 10126 |
|  | PDB: 6S6B | PDB: 6S8E | PDB: 6S8B | | PDB: 6S91 |
| **Data collection** |  | | | | |
| Electron Microscope | Titan Krios | | | | |
| Voltage (kV) | 300 | | | | |
| Electron detector | Falcon 3 | | | | |
| Electron dose (e-/Å^2^) | 40 (40 frms) | 85.5 (19 frms, LM) | | 40 (50 frms) | |
| Defocus range (μm) | -1.7 to -2.6 | | | | |
| Pixel size (Å) | 0.832 | | | | |
| **3D Reconstruction** |  | | | | |
| Raw images | 2,574 | 5,190 | 3,763 | | 3,763 |
| Initial particles | 614 K | 344 K | 785 K | | 785 K |
| Final particles | 66 K | 112 K | 82 K | | 52 K |
| Map resolution (Å) | 2.75 | 3.09 | 2.41 | | 2.68 |
| FSC threshold | 0.143 | 0.143 | 0.143 | | 0.143 |
| Map sharpening B-factor (Å^2^) | -63.5 | -110.3 | -55.0 | | -52.7 |
| **Model composition** |  | | | | |
| Protein residues | 8194 | 7502 | 7479 | | 7488 |
| Non-hydrogen atoms | 66914 | 62225 | 62153 | | 62051 |
| **Model refinement** |  | | | | |
| B factors (Å^2^) |  |  |  |  |  |
| Protein | 96.14 | 93.65 | 70.61 | | 59.04 |
| Nucleotide | 74.83 | 73.74 | 70.79 | | 55.16 |
| Ligand | 72.11 | 156.7 | 68.42 | | 57.92 |
| R.m.s. deviations |  | | | | |
| Bond length (Å) | 0.005 | 0.06 | 0.008 | | 0.006 |
| Bond angles (°) | 0.757 | 0.74 | 0.842 | | 0.775 |
| Ramachandran statistics (%) |  | | | | |
| Favored | 84.31 | 84.95 | 83.67 | | 86.29 |
| Allowed | 15.29 | 14.81 | 15.82 | | 13.55 |
| Outlier | 0.41 | 0.24 | 0.51 | | 0.16 |
| Rotamer outlier (%) | 0.15 | 0.16 | 0.26 | | 0.09 |
| MolProbity score | 2.43 | 2.42 | 2.45 | | 2.42 |
| Clash score | 18.31 | 18.27 | 18.76 | | 19.34 |
| Model vs Data CC (mask) | 0.82 | 0.79 | 0.78 | | 0.74 |

|  | Cmr1_1_2_1_3_1_4_4_5_3_6_1_7_22_β^crRNA^-CTR | Cmr1_1_2_1_3_1_4_4_5_3_6_1_7_26_β^crRNA^-CTR-AMPPNP, state1, polyT | Cmr1_1_2_1_3_1_4_4_5_3_6_1_7_26_β^crRNA^-CTR-AMPPNP, state2, polyT |
| --- | --- | --- | --- |
|  | EMDB: 10109 | EMDB: 10197 | EMDB: 10196 |
|  | PDB: 6SIC | PDB: 6SHB | PDB: 6SH8 |
| **Data collection** |  | | |
| Electron Microscope | Titan Krios | | |
| Voltage (kV) | 300 | | |
| Electron detector | Falcon 3 | | |
| Electron dose (e-/Å^2^) | 40 (40 frms) | 85.5 (19 frms, LM) | 40 (50 frms) |
| Defocus range (μm) | -1.7 to -2.6 | | |
| Pixel size (Å) | 0.832 | | |
| **3D Reconstruction** |  | | |
| Raw images | 2,574 | 5,190 | 3,763 |
| Initial particles | 614 K | 344 K | 785 K |
| Final particles | 66 K | 112 K | 82 K |
| Map resolution (Å) | 2.75 | 3.09 | 2.41 |
| FSC threshold | 0.143 | 0.143 | 0.143 |
| Map sharpening B-factor (Å^2^) | -63.5 | -110.3 | -55.0 |
| **Model composition** |  | | |
| Protein residues | 8194 | 7502 | 7479 |
| Non-hydrogen atoms | 66914 | 62225 | 62153 |
| **Model refinement** |  | | |
| B factors (Å^2^) |  | | |
| Protein | 96.14 | 93.65 | 70.61 |
| Nucleotide | 74.83 | 73.74 | 70.79 |
| Ligand | 72.11 | 156.7 | 68.42 |
| R.m.s. deviations |  | | |
| Bond length (Å) | 0.005 | 0.06 | 0.008 |
| Bond angles (°) | 0.757 | 0.74 | 0.842 |
| Ramachandran statistics (%) |  | | |
| Favored | 84.31 | 84.95 | 83.67 |
| Allowed | 15.29 | 14.81 | 15.82 |
| Outlier | 0.41 | 0.24 | 0.51 |
| Rotamer outlier (%) | 0.15 | 0.16 | 0.26 |
| MolProbity score | 2.43 | 2.42 | 2.45 |
| Clash score | 18.31 | 18.27 | 18.76 |
| Model vs Data CC (mask) | 0.82 | 0.79 | 0.78 |

**Table S2. Oligonucleotide Substrates Used in This Study. Related to Key Resources Table.**

| Type | Sequence 5’ to 3’ | Source |
| --- | --- | --- |
| CRISPR RNA template | TTCAAAGCTTAGATACCCTGGAGGGAAACCAGACTTAACACCA | *In vivo* transcription |
| RNA (CTR) | UGUUAAGUCUGGUUUCCCUCCAGGGUAUCUAAGCUUUGAAAAAAAA | IDT |
| RNA (NTR) | UGUUAAGUCUGGUUUCCCUCCAGGGUAUCUAAGCUUUGAACUUUCAATAA | IDT |
| RNA (naTR) | UGUUAAGUCUGGUUUCCCUCCAGGGUAUCUAAGCUUUGAA | IDT |
| RNA (5+T6) | UUGAAAAAAAA | IDT |
| RNA (11+T6) | UAAGCUUUGAAAAAAAA | IDT |
| RNA (17+T6) | GGUAUCUAAGCUUUGAAAAAAAA | IDT |
| RNA (23+T6) | CUCCAGGGUAUCUAAGCUUUGAAAAAAAA | IDT |
| RNA (29+T6) | GUUUCCCUCCAGGGUAUCUAAGCUUUGAAAAAAAA | IDT |
| RNA (34+T6) | GUCUGGUUUCCCUCCAGGGUAUCUAAGCUUUGAAAAAAAA | IDT |
| RNA (40+T1) | UGUUAAGUCUGGUUUCCCUCCAGGGUAUCUAAGCUUUGAAA | IDT |
| RNA (40+T2) | UGUUAAGUCUGGUUUCCCUCCAGGGUAUCUAAGCUUUGAAAA | IDT |
| RNA (40+T3) | UGUUAAGUCUGGUUUCCCUCCAGGGUAUCUAAGCUUUGAAAAA | IDT |
| RNA (40+T4) | UGUUAAGUCUGGUUUCCCUCCAGGGUAUCUAAGCUUUGAAAAAA | IDT |
| RNA (S10RNA） | AUAGAAUGCCCCCAUUAUACAAUAUCUACGUUUUAGAUGAAAAAAA | IDT |
| RNA  (RNA2) | FAM-CGUCUCUCUCCUGGAAUACAGAUGCUGGUGUUCGUGG | IDT |
| DNA (S10DNA) | ACTATAGGGAGAATAGAATGCCCCCATTATACAATATCTACGTTTTAGATGAACGCGTCT | IDT |
| DNA (polyT 20-mer) | TTTTTTTTTTTTTTTTTTTT | IDT |

**Table S3. DNA Primers Used for Construction of Plasmids. Related to Key Resources Table.**

| Oligonucleotides | Sequence 5’ to 3’ |
| --- | --- |
| F-NdeI | GCGTGACCATATGGCAATCGACTTTTTAGTC |
| R-mut | GGAATCGCAAATGGGTTTT |
| F-mut | AAAACCCATTTGCGATTCC |
| R-NotI | ATAAGAATGCGGCCGCTTTTATCACCTCTATCTTCC |
| MCS-fwd | ATGCCCCGGGATGTTAAACAAGTTAGG |
| MCS-rev | GGCACTCGAGAAAAAAAAGATTTTGCTTAATGGTG |
| 2β F_NheI | CTAGCTAGCATGAGCATTGATAATGTTCT |
| 2β 650D soeR | AAGTAATCGCTAGCAGCCCTAACTAAAGC |
| 2β 652D soeR | CTAAGTAAGCGCTATCAGCCCTAAC |
| 2β 650D soeF | GGGCTGCTAGCGATTACTTAGGAGATTTG |
| 2β 652D soeF | GATAGCGCTTACTTAGGAGATTTGTTAG |
| 2β R_ NotI | AAGGAAAAAAGCGGCCGCTCACTTCTCGCCACCGTAAAT |
| Del_2β L-F SalI | TTACGC GTCGAC TCTTTCGATCCTTATATGAACTTGC |
| Del_2β SOE-R | GAAATTAGATCATTAGGAATTGCAATAATTTTGTAGTCTA |
| Del_2β SOE-F | TAGACTACAAAATTATTGCAATTCCTAATGATCTAATTTC |
| Del_2β R-R NotI | AAGGAAAAAAGCGGCCGCTATCATAAAAAAGTTCATAAATCCC |
| Del_2β spacer F | AAAGTTAGATCTTCACAGATTTCGTTTTTCTTCTCAATCATTTC |
| Del_2β spacer R | TAGCGAAATGATTGAGAAGAAAAACGAAATCTGTGAAGATCTAA |
| GE_2βL-F SalI | TTACGC GTCGAC TCTTTCGATCCTTATATGAACTTGC |
| GE_2βHD SOE-R | CCCATGCCTTATTTGGAGGAGCAGCAAGTAATGCAAT |
| GE_2βHD SOE-F | ATAAGGCATGGG TAATAACAGGTCG |
| GE_2βRHD R-R NotI | AAGGAAAAAAGCGGCCGCATGGTCAAAAATTGTGTGAGTAGGG |
| GE_2βHD spacer-F | AAAGGAGGATCATGAAGTAATGCAATAATTTTGTAGTCTAAGAA |
| GE_2βHD spacer-R | TAGCTTCTTAGACTACAAAATTATTGCATTACTTCATGATCCTC |
| GE_2βGD L-F SalI | TTACGC GTCGACGTTAGGGCTGATAGCGATTA |
| GE_2βGD SOE-R | CATTGCCAATAATGCAGCAGCTGCCGCATAAATTAC |
| GE_2βGD SOE-F | GTAATTTATGCGGCAGCTGCTGCATTATTGGCAATG |
| GE_2βRGD R-R NotI | AAGGAAAAAAGCGGCCGCAATCGTAAAGTAAGGACACA |
| GE_2βGD spacer-F | AAAGCCCCGCATAAATTACAAATCCCTTATGTTTATTAACCAAT |
| GE_2βGD spacer-R | TAGCATTGGTTAATAAACATAAGGGATTTGTAATTTATGCGGGG |
| GE_4β D31A L-F SalI | TTACGCGTCGACTTAATTATGCTGGGAATCTTATTGT |
| GE_4βD31A SOE-R | GCCTAGTTCATCCCTTTGGAATGGTAATGCTATTTCCTC |
| GE_4βD31A SOE-F | GAGGAAATAGCATTACCATTCCAAAGGGATGAACTAGGC |
| GE_4βD31A R-R NotI | AAGGAAAAAAGCGGCCGCTAGATCCTTGTGATTATGTTAAAGG |
| GE_4βD31A spacer-F | GAAAGGGTAAGTCTATTTCCTCCTCTACAGATGAGCCAGA |
| GE_4βD31A spacer-R | TCTGGCTCATCTGTAGAGGAGGAAATAGACTTACCCTTTC |
| Del_7βL-F SalI | TTACGCGTCGACTAGGGGTTGGTCACGCCCACTTCCG |
| Del_7βR-R NotI | AAGGAAAAAAGCGGCCGCATTGCTGAAGGAAGCGTTAG |
| Del_7β SOE-R | GTCATCTGATATATTCCCCTTAATACTATTTGTAATCGGTA |
| Del_7β SOE-F | TACCGATTACAAATAGTATTAAGGGGAATATATCAGATGAC |
| Del_7β spacer F | AAAGGTGATTACTAGAACATTATGAATGCCTCTCTCATCTATAT |
| Del_7β spacer R | TAGCATATAGATGAGAGAGGCATTCATAATGTTCTAGTAATCAC |
