## Supplementary material for "Structures of the Cmr-β Complex Reveal the Regulation of the Immunity Mechanism of Type III-B CRISPR-Cas": supplemntary figs 1-8

**Fig. S1**

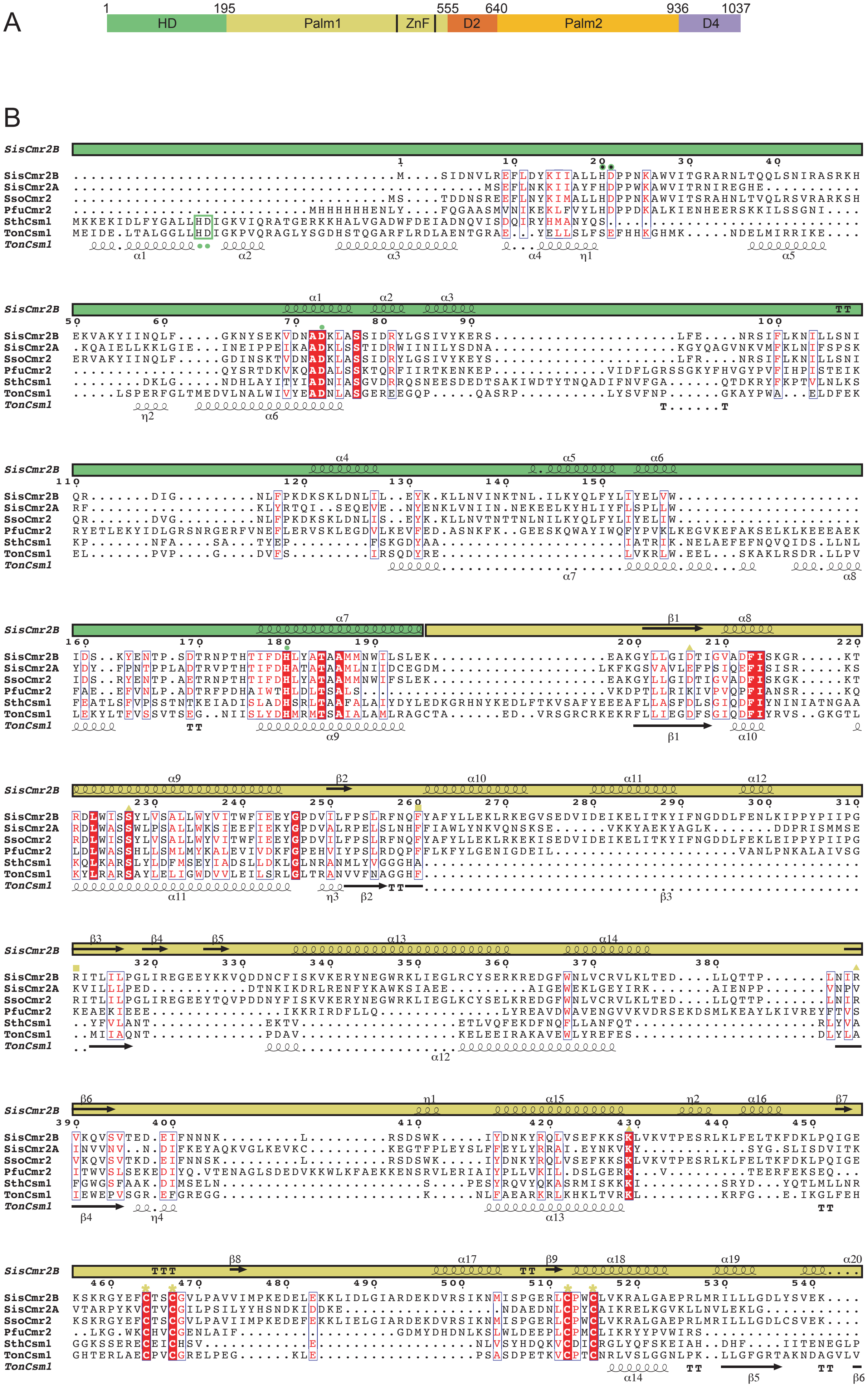
A
1
195
555
640
936
1037
HD
Palm1
ZnF
D2
Palm2
D4
B
SisCmr2B
SisCmr2A
SsoCmr2
PfuCmr2
SthCsm1
TonCsm1

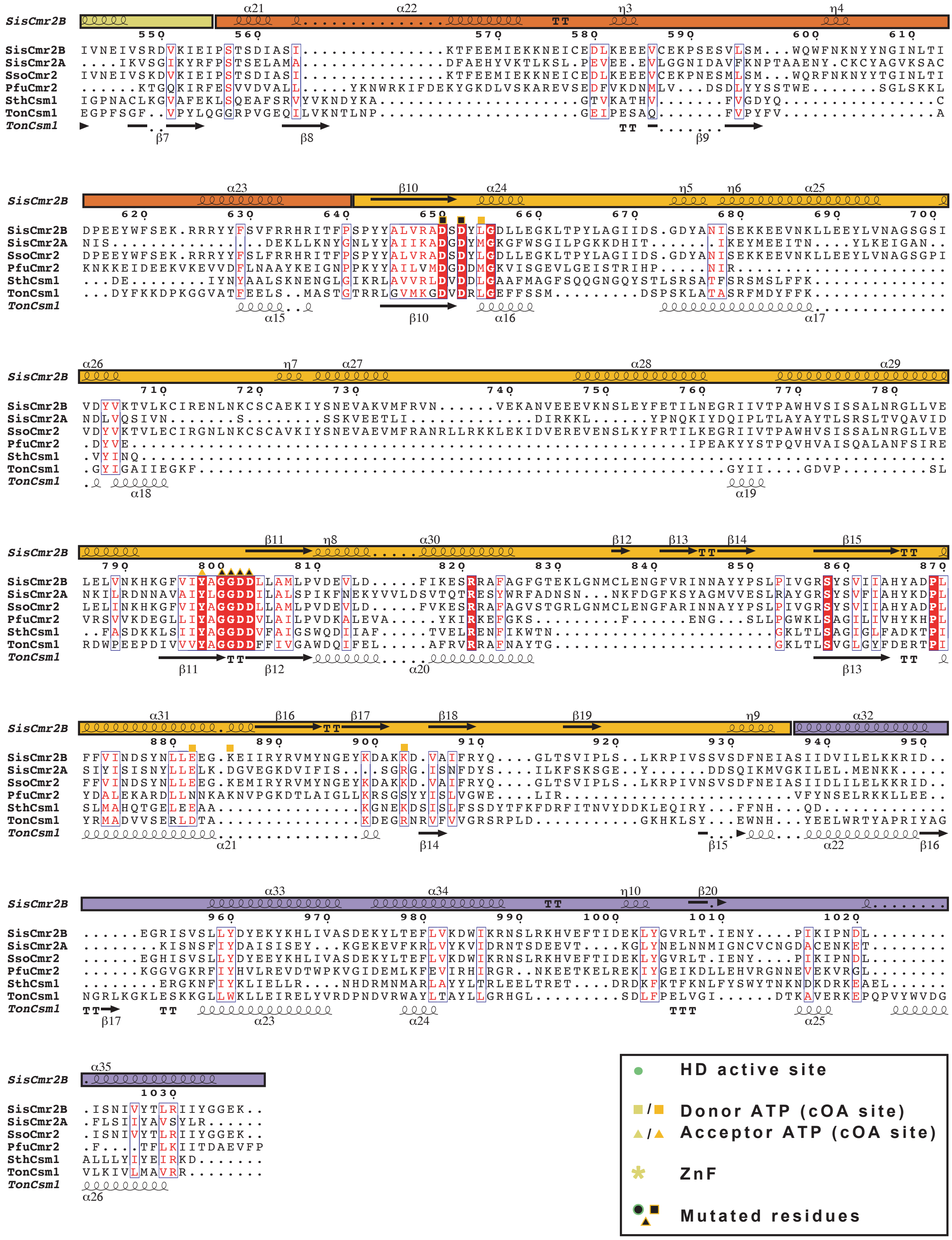
SisCmr2B IVNEIVSRDVKIEIPSTSDIASI...............KTFEEMIEKKNEICEDLKEEEVCEKPSESVLSM...WQWFNKNYYNGINLTI
SisCmr2A ....IKVSGIKYRFPSTSELAMA...............DFAEHYVKTLKSL.PEVEE.EVLGGNIDAVFKNPTAAENY.CKCYAGVKSAC
SsoCmr2 IVNEIVSKDVKIEIPSTSDIASI...............KTFEEMIEKKNEICEDLKEEEVCEKPNESMLSM...WQRFNKNYYTGINLTI
PfuCmr2 ......KTGQKIRFESVVDVALL.....YKNWRKIFDEKYGKDLVSKAREVSEDFVKDNMLV...DSDLYYSSTWE.......SGLSKKL
SthCsm1 IGPNACLKGVAFEKLSQEAFSRVYVKNDYKA.....................GTVKATHV.......FVGDYQ................C
TonCsm1 EGPFSGF..VPYLQGGRPVGEQILVKNTLNP.....................GEIPESAQ.......FVPYFV................A
SisCmr2B DPEEYWFSEK.RRRYYFSVFRRHRITFPSPYYALVRADSDYLGDLLEGKLTPYLAGIIDS.GDYANISEKKEEVNKLLEEYLVNAGSGSI
SisCmr2A NIS..............DEKLLKNYGNLYYAIIKADGDYMGKGFWSGILPGKKDHIT.......IKEYMEEITN....YLKEIGAN..
SsoCmr2 DPEEYWFSEK.RRRYYFSLFRRHRITFPSPYYALVRADSDYLGDLLEGKLTPYLAGIIDS.GDYANISEKKEEVNKLLEEYLVNAGSGPI
PfuCmr2 KNKKEIDEEKVKEVVDFLNAAYKEIGNPPKYYAILVMDGDDMGKVISGEVLGEISTRIHP.....NIR.....................
SthCsm1 ...DE........IYNYAALSKNENGLGIKRLAVVRLDVDDLGAAFMAGFSQQGNGQYSTLSRSATFSRSMSLFFK.............
TonCsm1 ...DYFKKDPKGGVATFEELS..MASTGTRRLGVMKGDVDRLGEFFSSM.........DSPSKLATASRFMDYFFK.............
SisCmr2B VDYVKTVLKCIRENLNKCSCAEKIYSNEVAKVMFRVN......VEKANVEEEVKNSLEYFETILNEGRIIVTPAWHVSISSALNRGLLVE
SisCmr2A NDLVQSIVN.......S........SSKVEETLI............DIRKKL......YPNQKIYDQIPLTLAYAYTLSRSLTVQAVID
SsoCmr2 VDYVKTVLECIRGNLNKCSCAVKIYSNEVAEVMFRANRLLRKKLEKIDVEREVENSLKYFRTILKEGRIIVTPAWHVSISSALNRGLLVE
PfuCmr2 .DYVE.....................................................IPEAKYYSTPQVHVAISQALANFSIRE
SthCsm1 .VYINQ.....................................................................
TonCsm1 .GYIGAIIEGKF.............................................GYII....GDVP.........SL
SisCmr2B LELVNKHKGFVIYAGGDDLLAMLPVDEVLD.....FIKESRRAFAGFGTEKLGNMCLENGFVRINNAYYPSLPIVGRSYSVIIAHYADPL
SisCmr2A NKILRDNNAVAIYLGGDDILALSPIKFNEKYVVLDSVTQTRESYWRFADNSN...NKFDGFKSYAGMVVESLRAYGRSYSVFIAHYKDPL
SsoCmr2 LELINKHKGFVIYAGGDDLLAMLPVDEVLD.....FVKESRRAFAGVSTGRLGNMCLENGFARINNAYYPSLPIVGRSYSVIIAHYADPL
PfuCmr2 VRSVVKDEGLLIYAGGDDVLAILPVDKALEVA.....YKIRKEFGKS.........F......ENG..SLLPGWKLSAGILIVHYKHPL
SthCsm1 ..FASDKKLSIIYAGGDDVFAIGSWQDIIAF.....TVELRENFIKWTN.......................GKLTLSAGIGLFADKTPI
TonCsm1 RDWPEEPDIVVVYAGGDDFFIVGAWDQIFEL.....AFRVRRAFNAYTG..........................GKLTLSVGLGYFDERTPI
SisCmr2B FFVINDSYNLLEEG.KEIIRYRVMYNGEYKDAKKD.VAIFRYQ....GLTSVIPLS..LKRPIVSSVSDFNEIASIIDVILELKKRID..
SisCmr2A SIYISISNYLLELK.DGVEGKDVIFIS...SGRG.ISNFDYS....ILKFSKSGE..Y......DDSQIKMVGKILEL.MENKK.....
SsoCmr2 FFVINDSYNLLEEG.KEMIRYRVMYNGEYKDAKKD.VAIFRYQ....GLTSVIPLS..LKRPIVNSVSDFNEIASIIDLILELKKRID..
PfuCmr2 YDALEKARDLLNNAKNVPGKDTLAIGLLKRSGSYYISLVGWE....LIR.........................VFYNSELRKKLLEE..
SthCsm1 SLMAHQTGELEEAA..............KGNEKDSISLFSSDYTFKFDRFITNVYDDKLEQIRY...FFNH...QD......
TonCsm1 YRMADVVSERLDTA..............KDEGRNRVFVVGRSRPLD.........GKHKLSY...EWNH...YEELWRTYAPRIYAG
SisCmr2B ......EGRISVSLLYDYEKYKHLIVASDEKYLTEFLVKDWIKRNSLRKHVEFTIDEKLYGVRLT.IENY....PIKIPNDL........
SisCmr2A ........KISNSFIYDAISISEY....KGEKEVFKRLVYKVIDRNTSDEEVT....KGLYNELNNMIGNCVCNGDACENKET.......
SsoCmr2 ......EGHISVSLLYDYEEYKHLIVASDEKYLTEFLVKDWIKRNSLRKHVEFTIDEKLYGVRLT.IENY....PIKIPNDL........
PfuCmr2 ......KGGVGKRFIYHVLREVDTWPKVGIDEMLKFEVIRHIRGR..NKEETKELREKIYGEIKDLLEHVRGNNEVEKVRGL........
SthCsm1 ........ERGKNFIYKLIELLR....NHDRMNMARLAYYLTRLEELTRET....DRDKFKTFKNLFYSWYTNKNDKDRKEAEL......
TonCsm1 NGRLKGKLESKKGLLWKLLEIRELYVRDPNDVRWAYLTAYLLGRHGL.........SDLFPELVGI......DTKAVERKEPQPVYWVDG
SisCmr2B .ISNIVYTLRIIYGGEK..
SisCmr2A .FLSIIYAVSYLR......
SsoCmr2 .ISNIVYTLRIIYGGEK..
PfuCmr2 .F....TFLKIITDAEVFP
SthCsm1 ALLLYIYEIRKD.......
TonCsm1 VLKIVLMAVRR........
HD active site
Donor ATP (cOA site)
Acceptor ATP (cOA site)
ZnF
Mutated residues

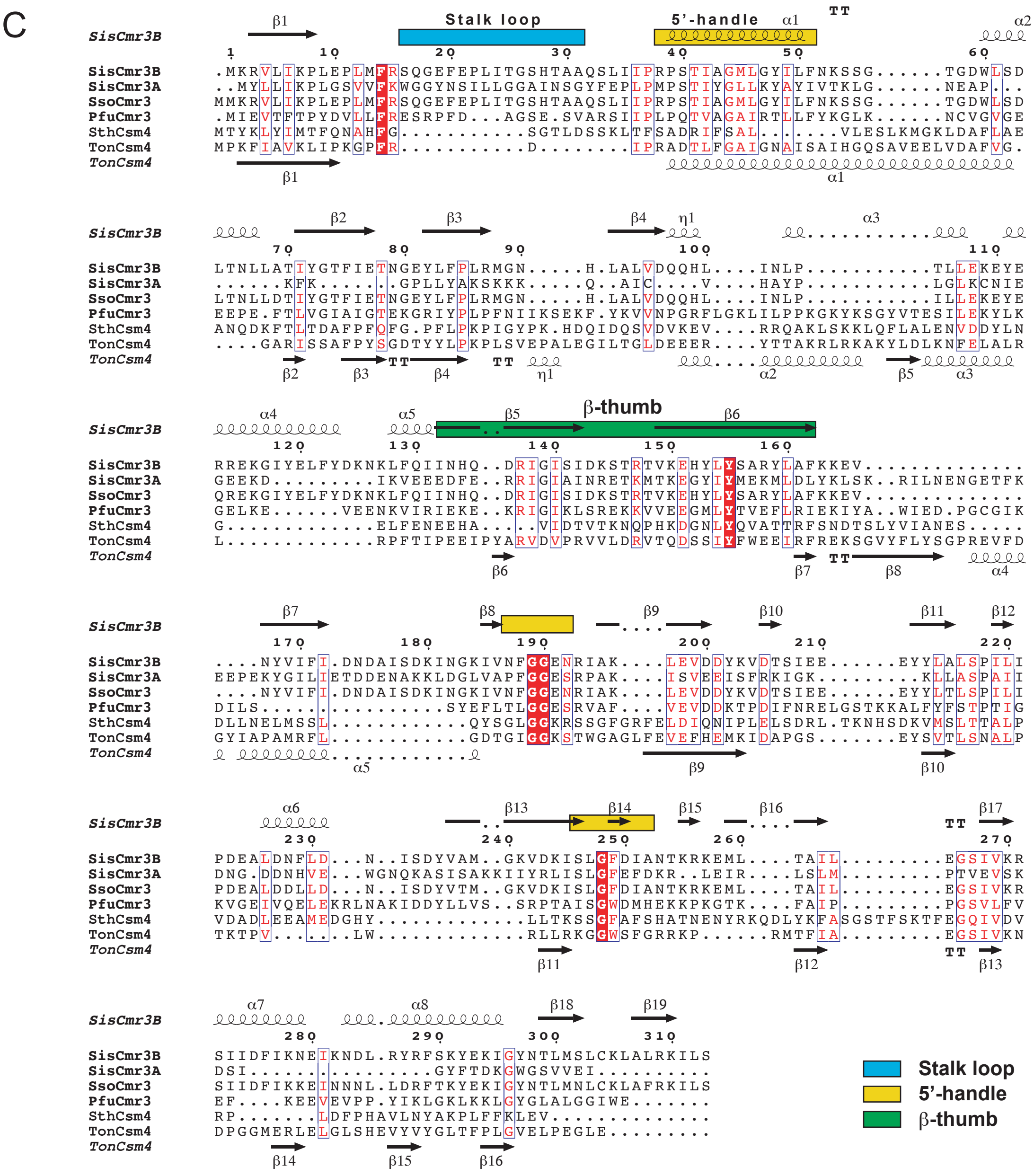

**Figure S1. Sequence alignments of the Cmr2 and Cmr3 subunits, related to Figure 1-4.**
(A) Schematic showing the domain borders and composition of the SisCmr2 protein.
(B) Sequence alignment of SisCmr2B against type IIIA representatives SisCmr2A, SsoCmr2, PfuCmr2, and Type IIIA representatives TonCsm1 and SthCsm1. The Cmr2 domains and secondary structure indicated together above the aligned sequences. Residues involved in the HD active center, coordinating to either the donor or acceptor AMPPnPs (ATPs), ligating the ZnF zinc ion, and residues mutated in the presented work, are all indicated according to the legend in the bottom right corner.
(C) Sequence alignment of SisCmr3B against Type IIIB representatives SisCmr3A, SsoCmr3, PfuCmr3, and Type IIIA representatives TonCsm4 and SthCsm4. Stalk loop, 5'-handle, and beta-thumb, are indicated above the aligned sequences (colored as indicated in the bottom right corner). The alignments were performed with ClustalO and rendered using ESPript 3.0.

Fig. S2

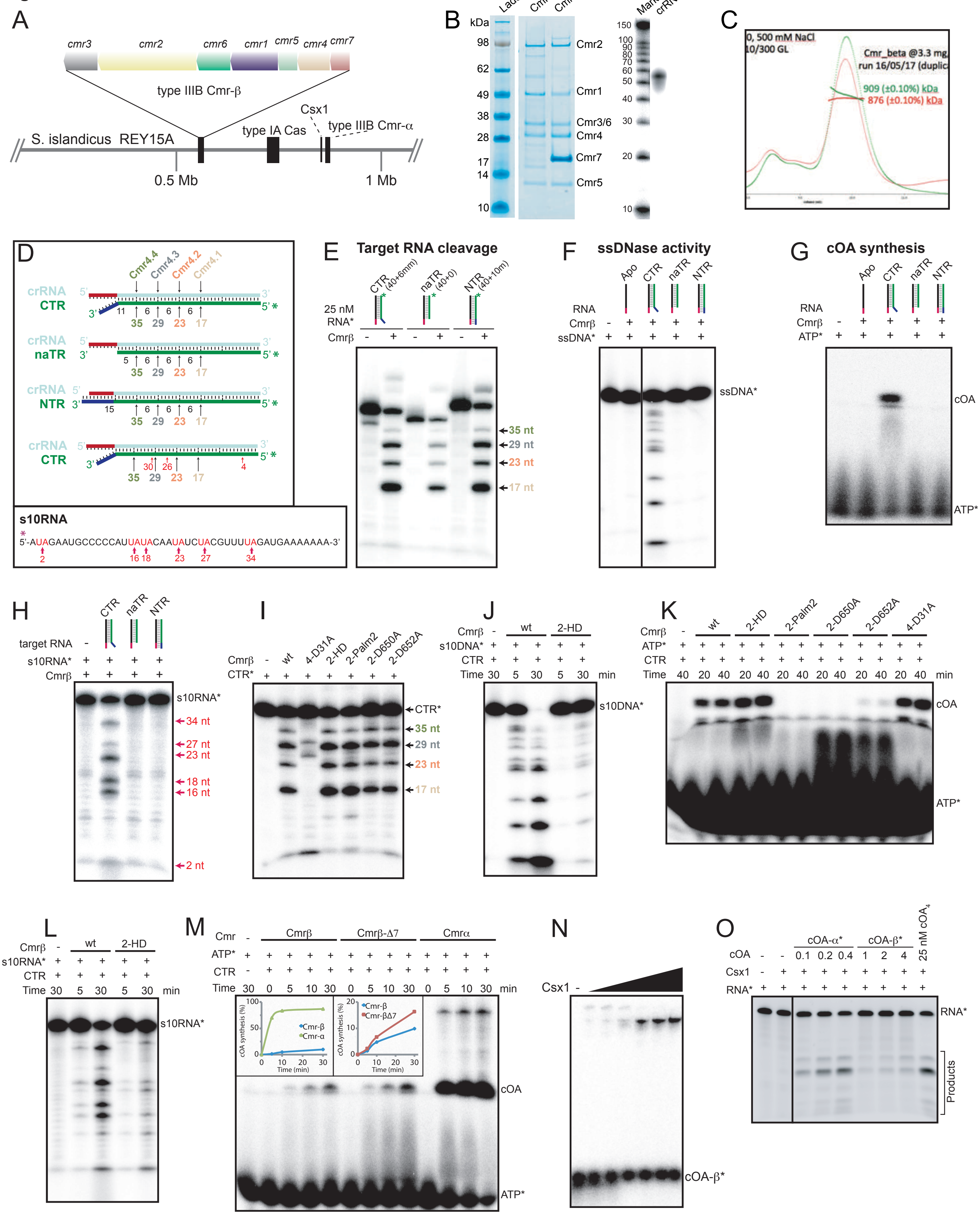


**Figure S2. Purification and Activities of the SisCmrB Complex, Related to Figure 1-4**

(A) Location of the type III-B CmrB gene locus.

(B) SDS-PAGE gel showing the purified CmrB complex, and an urea gel showing an RNA length marker and the crRNA extracted from the CmrB complex.

(C) Chromatogram of the MALS profile of purified CmrB complex.

(D) Schematic showing the crRNA hybridized with the different target RNAs used, and the cleavage positions resulting in the experimentally observed fragments. The bottom panel shows the sequence and UA cleavage sites of the s10RNA.

(E) Target RNA cleavage by CmrB in the presence and absence of CTR, naTR, and NTR, respectively. The four 5'-labelled fragments produced are indicated, with the color code used corresponding to the color of the Cmr4 subunit in the schematic (Fig 1C) which cleaves at the associated nucleotide position.

(F) and (G) ssDNA cleavage and cOA synthesis, respectively, by CmrB in the presence of CTR, naTR, and NTR.

(H) UA RNA cleavage by CmrB in the presence of high concentrations (525 nM) of CTR.

(I) Cognate target RNA cleavage by various mutants.

(J) ssDNA cleavage of the s10DNA substrate by the wt and HD-mutant complexes, at two time points.

(K) cOA synthesis in the presence of CTR and ATP, by wt and various Cmr2 mutant complexes.

(L) ssRNA cleavage of the s10RNA substrate by the wt and HD-mutant complexes, at two time points.

(M) cOA synthesis by the wt-beta, delta7, and wt-alfa complexes, in the presence of CTR and ATP. The inserts show the quantifications of the bands.

(N) Band shift upon Csx1 binding of the cOA produced by CmrB.

(O) RNA cleavage by Csx1 upon activation by cOA derived from either the alfa or beta complex reaction mixtures (exact concentration unknown), of 25 nM purified material from the CmrA reaction.

FigS3

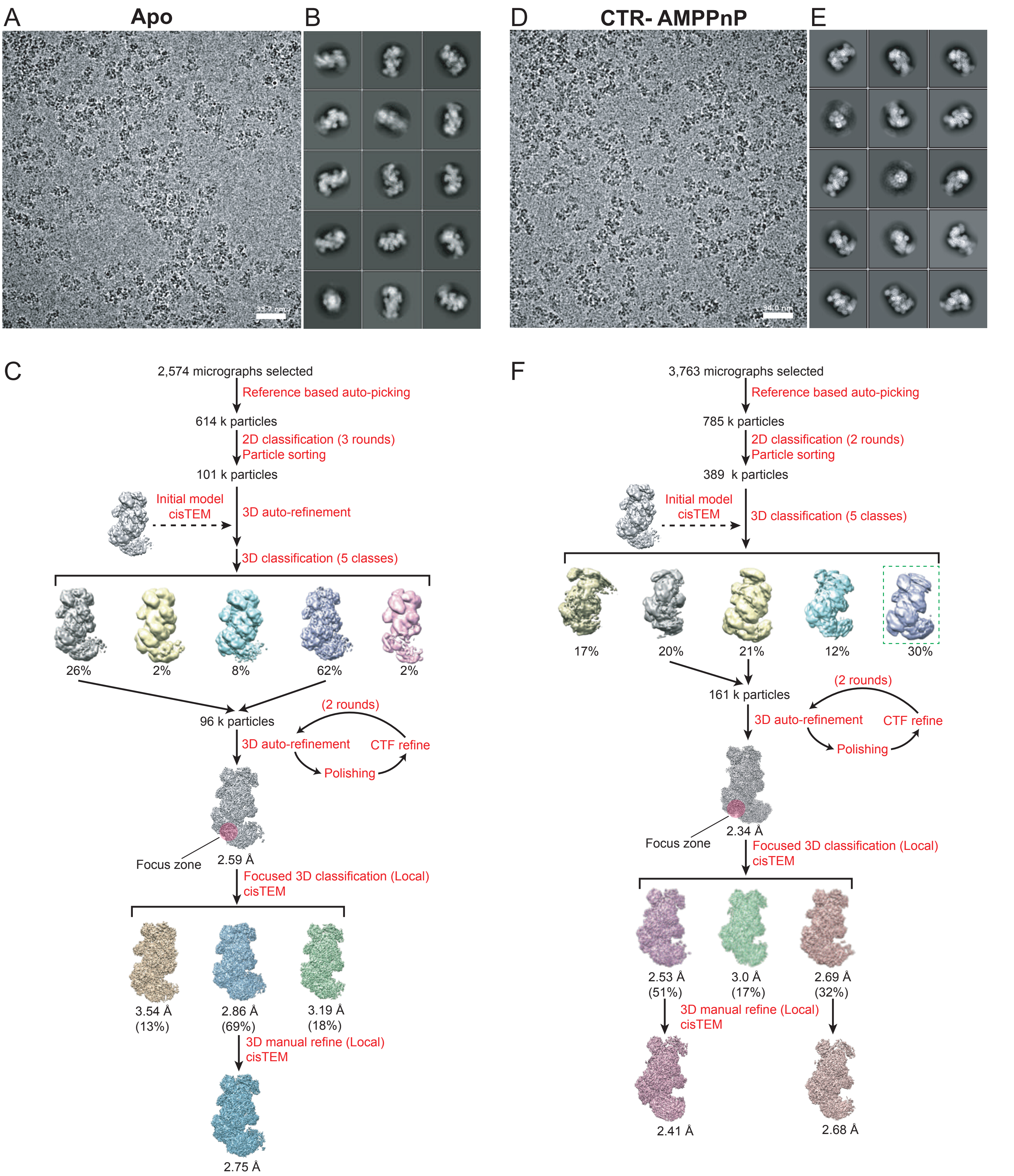


**Figure S3. Single Particle Cryo-EM Analysis of the Cmr-beta Apo and CTR-AMPPnP bound complexes, Related to Figure 1-4**

(A) and (D) Representative cryo-EM micrographs of the Cmr-beta apo and CTR-AMPPnP bound complexes, respectively, in vitreous ice on UltrAuFoil 0.6/1.0 grids.

(B) and (E) Reference-free 2D class averages sorted by class distribution (top-left to bottom-right).

(C) and (F) Overview of the cryo-EM data processing workflows for the Cmr-beta apo and CTR-AMPPnP bound complexes, respectively. The processing was generally performed in RELION-3.0. Steps performed in cisTEM are indicated.

FigS4

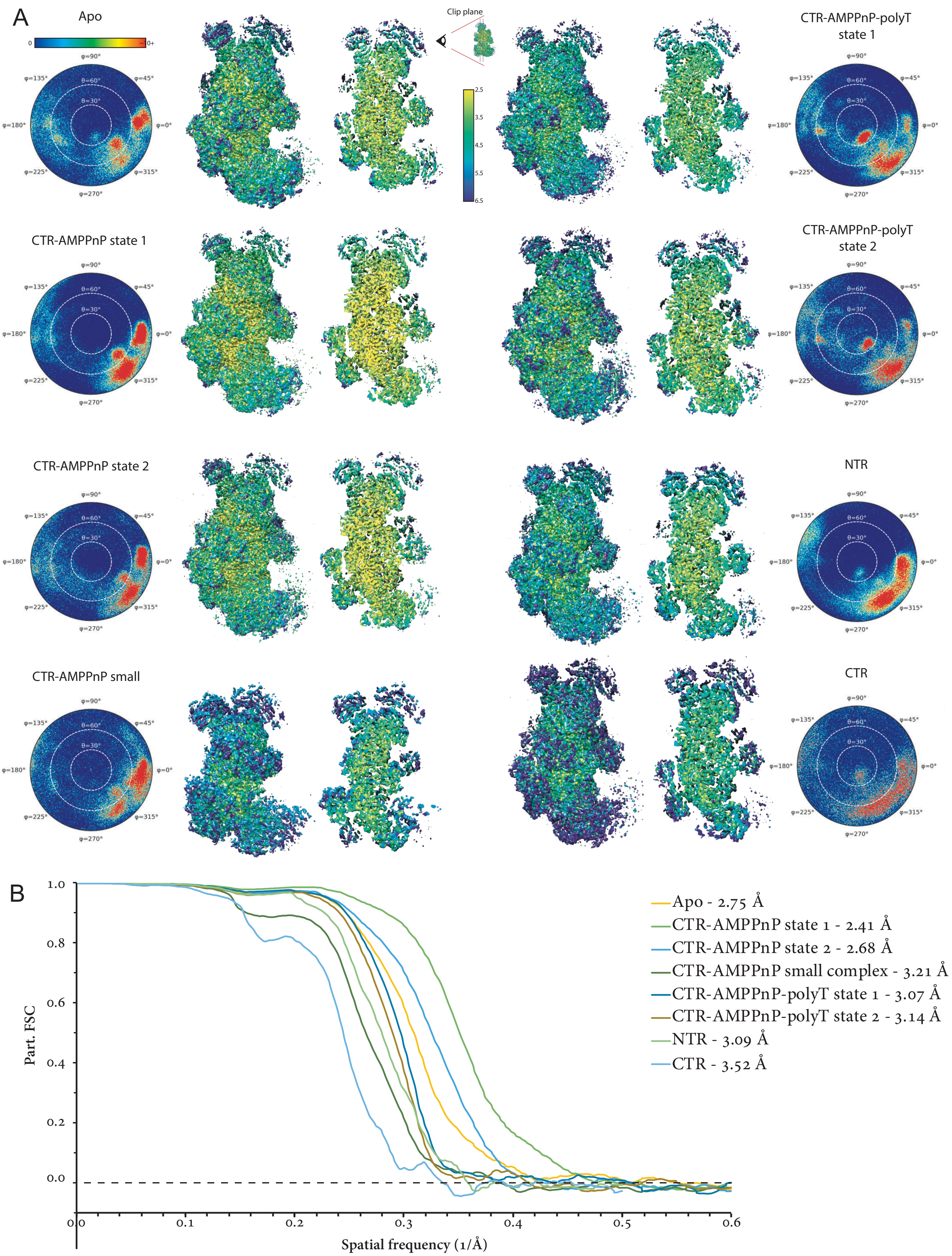


**Figure S4. Resolution Assesment and Validation of all Cryo-EM Density Maps, Related to Figure 1-3**
(A) Euler angle distribution plots showing the relative orientation of particles in the final 3D reconstructions, and local resolution maps and central slice through the maps.
(B) Fourier shell correlation (FSC) curves of the final 3D reconstructions.

**FigS5**

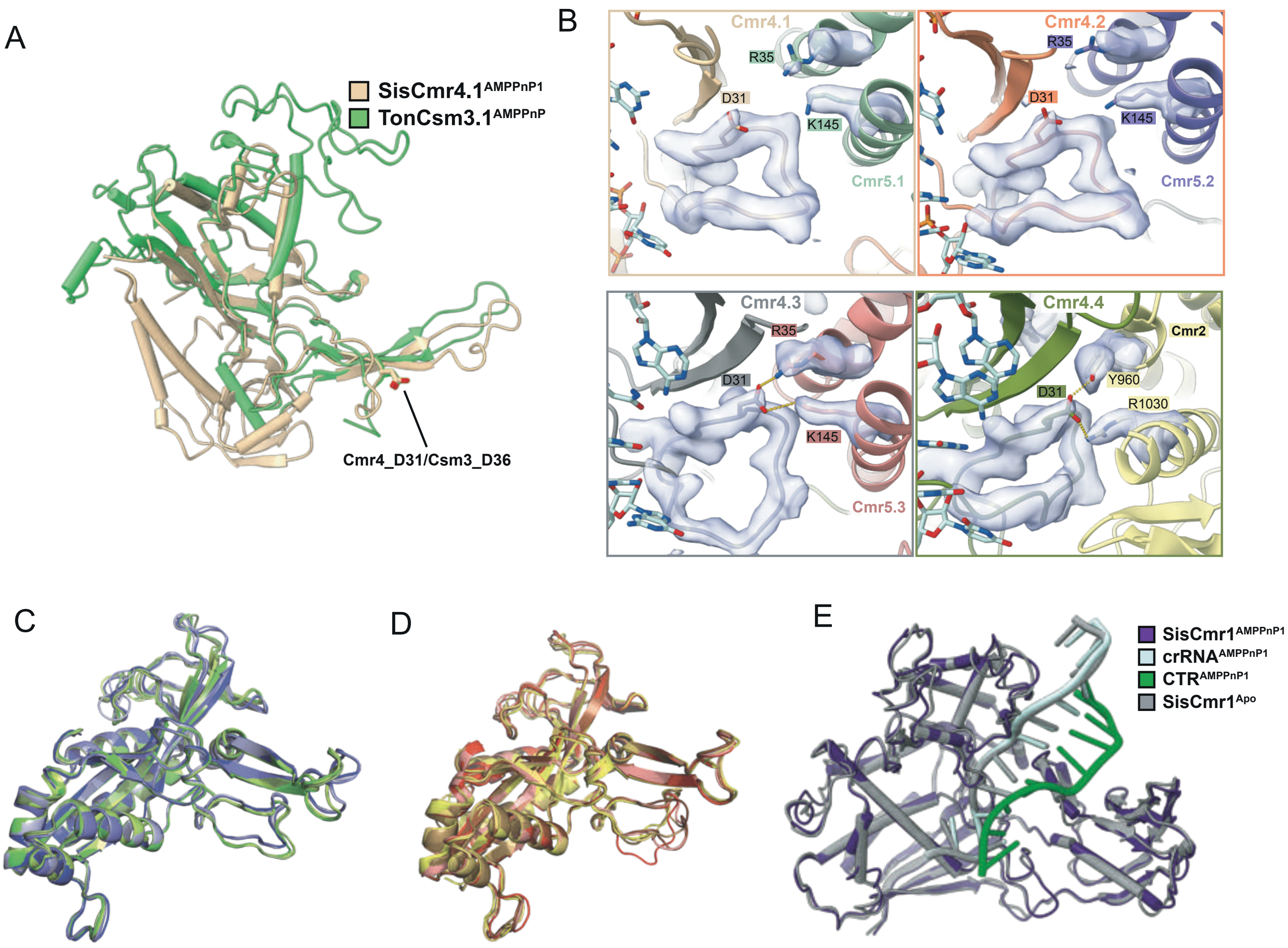


**Figure S5. Structural comparisons of SisCmrB, related to Figure 1-4.**
(A) Superposition of SisCmr4 with TonCsm3 (PDB 6O7E). The active site loop, beta-thumb, and a central part of the Cmr4/Csm3 proteins are structurally conserved, but only 57 residues were aligned overall, with an RSMD of 1.09 Å (22% sequence identity).
(B) Superposition of the Cmr4.1 subunits of the Apo and CTR-AMPPnP state 1 structures. Despite binding of the CTR, the RMSD is 0.642 covering 462 out of 476 residues.
(C) and (D) Superposition of the four Cmr4 subunits of the Cmr-β apo and Cmr-β-CTR-AMPPnP (class1) complexes, respectively. The AS loop (G21-F34) exhibits the greatest conformational changes. The Cmr4.2 and Cmr4.3 AS loops are compared in the close-up views.
(E) and(F) Comparison of the Cmr4 AS loops of the apo against the CTR-AMPPnP (class1) bound structures. Cmr4.3 (E) and Cmr4.4 (F) were selected as they exhibit the greatest displacement.

FigS6

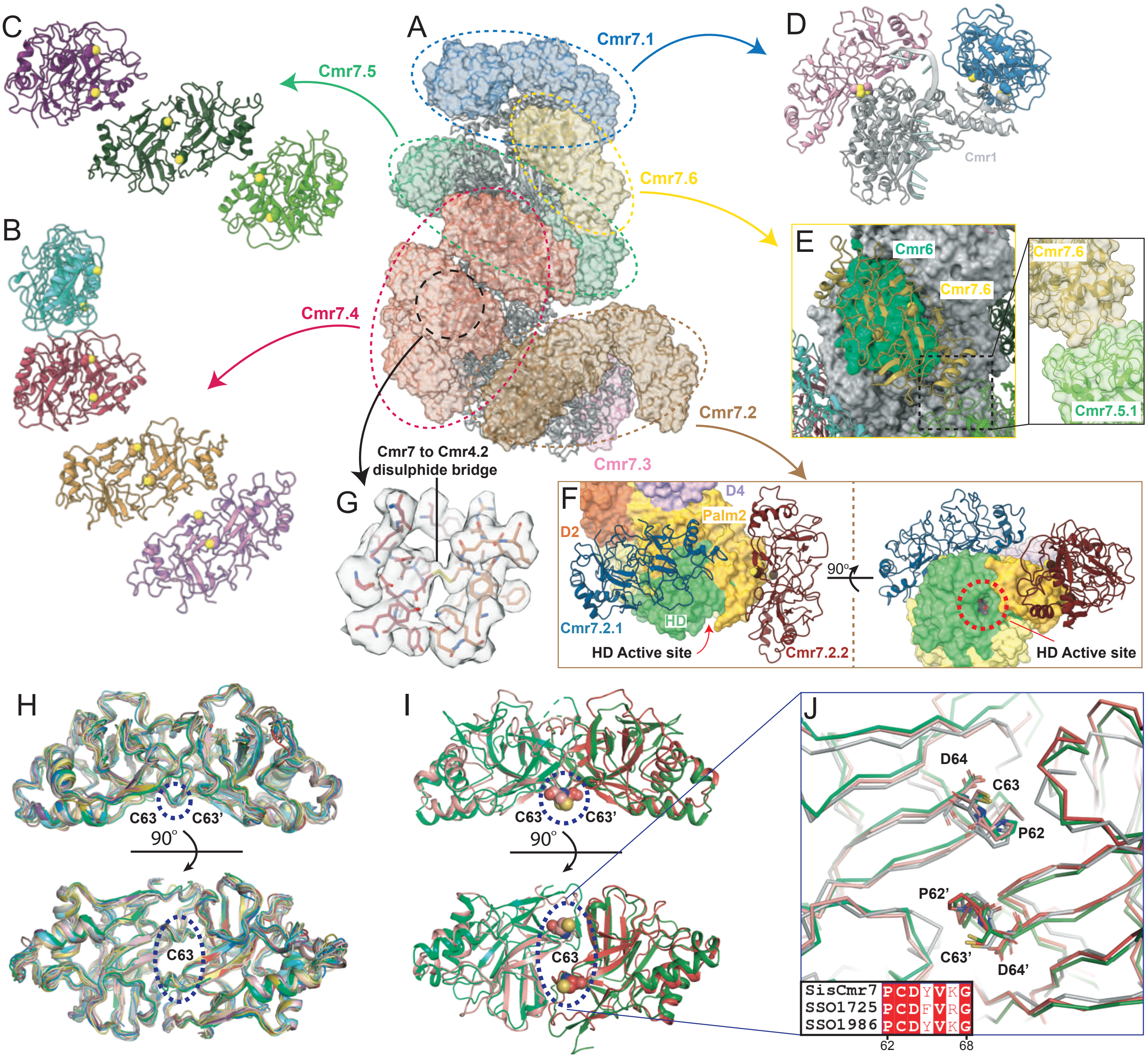


**Figure S6. Binding of Cmr7 dimers to the core complex, related to figure 1.**
Each Cmr7 dimer and each core subunit contributes one cysteine to a disulfide bridge linking the two units. The Cmr7 dimer has two centrally positioned cysteines. Cmr3, Cmr4, and Cmr5 only have one available cysteine, whereas Cmr6 has 2 cysteines. Cmr1 and Cmr2 have 11 and 13 cysteines, respectively.
(A) Overview showing the localization of the various Cmr7 dimers decorating the CmrB apo core complex (surface and cartoon, respectively). Cmr7 dimers are colored according to the identity of the subunit they bind.
(B) Four Cmr7 dimers are bound to the core complex via Cmr4 (red surface). The Cmr7.4 dimers stack upon each other in a helical arrangement. The Cmr7 cysteines available for disulfide bridge formation are shown in sphere representation.
(C) Three Cmr7 dimers are bound to the core complex via Cmr5 (green surface). The Cmr7.5 dimers interact in a different manner compared to Cmr7.4, although still in a regular pattern.
(D) Two Cmr7 dimers are bound to Cmr1 (blue surface). They do not interact with each other.
(E) One Cmr7 dimer is bound to Cmr6 sharing the largest interface area of any Cmr7 dimers bound to the core complex (1113 Å2) (yellow surface, cartoon). This dimer traverses the target RNA binding cleft and interacts with Cmr7.5.1, as shown in the inset, and is further stabilized thorugh interactions with Cmr1 (582 Å2).
(F) Two Cmr7 dimers are bound to Cmr2 (brown surface). Inset shows Cmr7 dimers in cartoon representation in two orientations, binding via the Palm2 domain (yellow surface), and wrapping around the HD domain (green surface). The location of the HD active site is indicated. Although Cmr2 cysteines 721 and 836 are located close to the cysteines of the Cmr7.2.2 and Cmr7.2.1 dimers, respectively, these disulfide bridges could not be modelled convincingly.
(G) Close-up view of the disulfide bridge and residues within 5 Å linking the Cmr7.4.2 dimer and Cmr4, and the density map of this region.
(H) Superimposition of all Cmr7 dimers from the Apo complex. The green dashed circle indicates the position of Cys63.
(I) Superimposition of all Cmr7 homologues from S. solfataricus (PDB 2XVO in grey, PDB 2X5Q in green) with Cmr7.4.2. RMSD of 4.17 Å and 1.91 Å, respectively. The superimposition of 2X5Q in green is visibly better, with most secondary structural elements conserved. (J) Close up view of the conserved PCD residues (in sticks) located on the center of the concave binding surface. An insertion into figure S6J shows a highly conserved portion of a sequence alignment of Cmr7 from S. islandicus (REY15A) and S. solfataricus (Sso1986 and Sso1725), at the PCD motif (residues 62-68).

**FigS7**

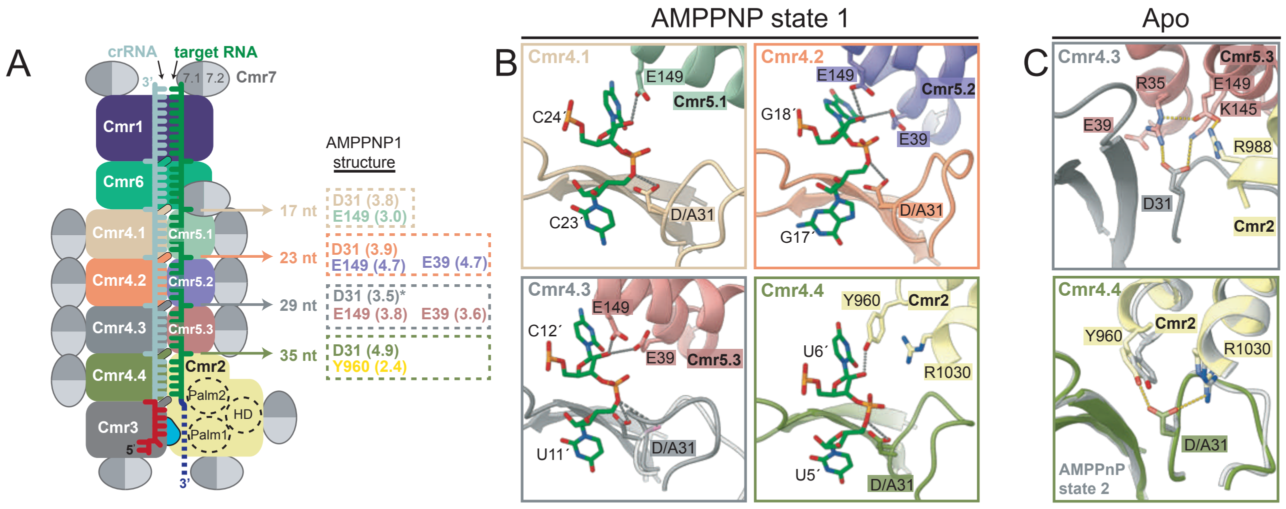


**Figure S7. Target RNA cleavage at the four Cmr4 active centers.**
(A) Overview schematic showing the origin of the 4 target RNA cleavage fragments - 17, 23, 29, 35 nt respectively - observed when providing the complex with a 5'-labelled target RNA. Dashed boxes show the distances of the active site residues in the four Cmr4 active sites with respect to the functional group they target (AMPPnP state 1 structure), color coded according to the subunit they belong.
(B) Close-up views of the Cmr4 active sites of the AMPPnP1 structure, color coded according to the schematic in (G).
(C) The Cmr4 active sites of the bottom two subunits (Cmr4.3 and Cmr4.4) in the absence of bound target RNA.

**FigS8**

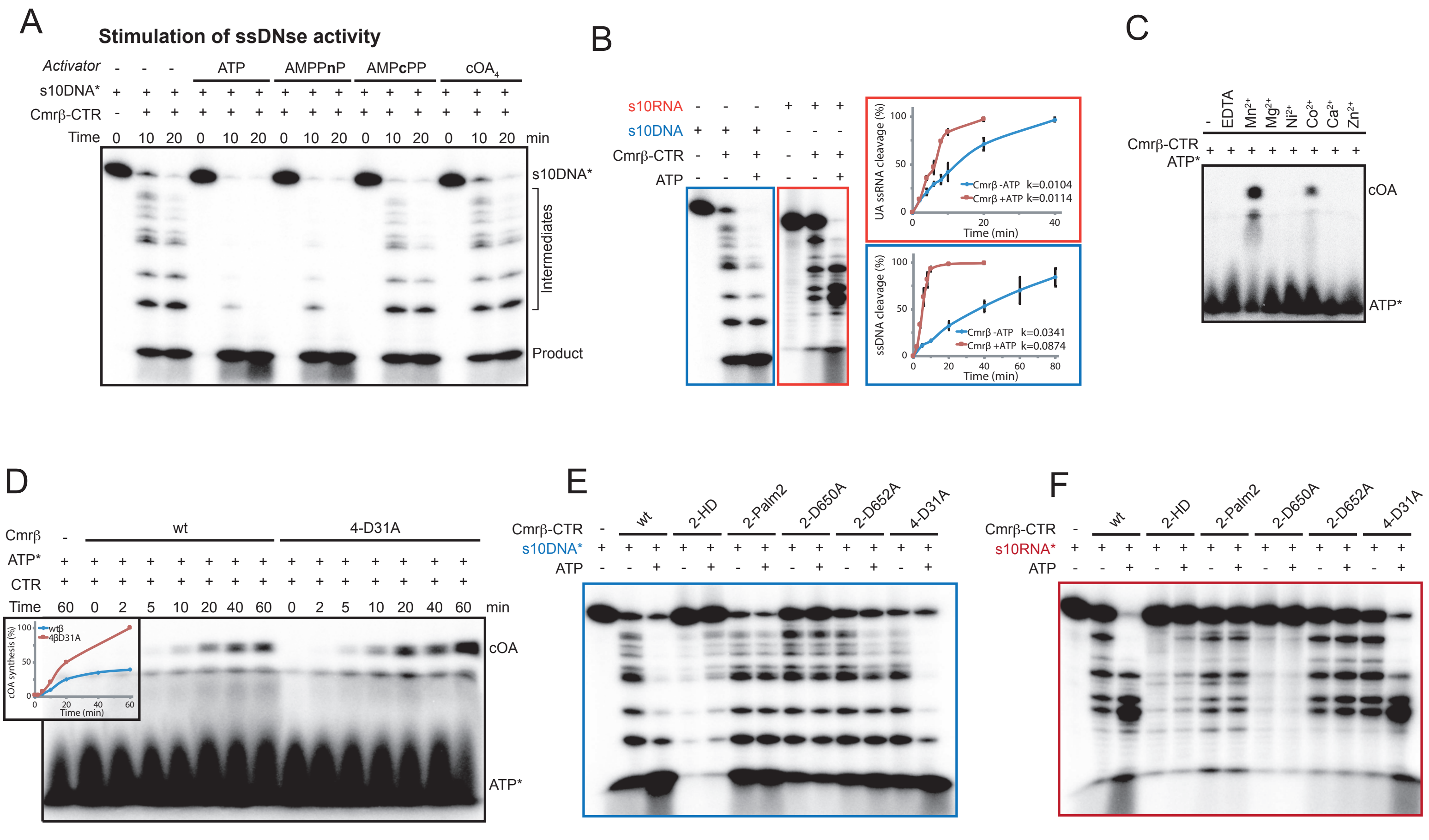


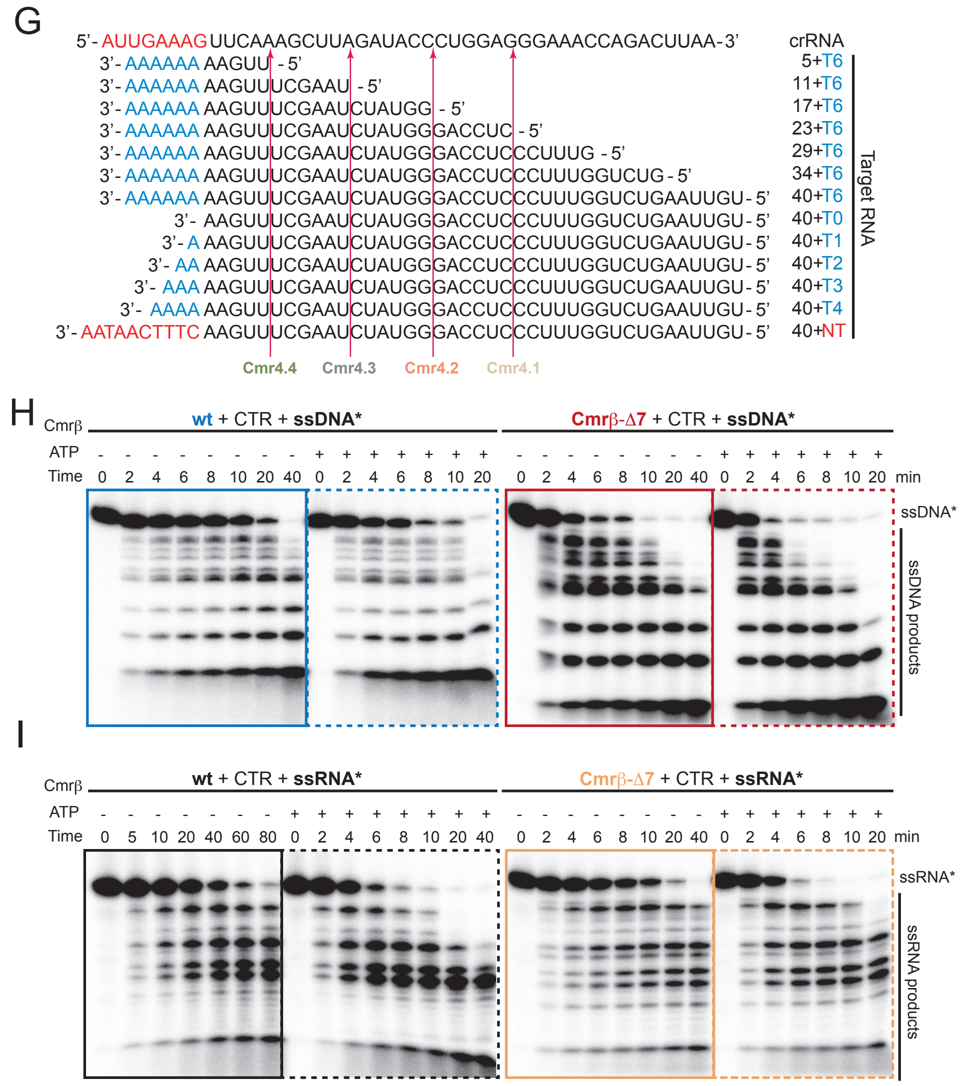


**Figure S8. Deciphering the role of Cmr7 and key residues on Cmr2 activities.**

(A) UA ssRNA (red) and ssDNA (blue) cleavage experiments using the s10RNA and s10DNA substrates respectively, and the CTR-activated CmrB in the presence and absence of ATP. The gels shown are single point experiments, whereas the quantification plots are based on time course experiments.

(B) cOA synthesis experiment in the presence of different divalent cations.

(C) cOA synthesis time course experiment in the presence of CTR, ATP, and either the wt or Cmr4 D31A mutant CmrB complexes. The insert is a quantification plot based on the gel.

(D) and (E) ssDNA and ssRNA cleavage experiments, respectively, in the presence and absence of ATP, comparing the wt CmrB complex with the mutants.

(F) The various target RNA sequences, including the short RNAs used in Fig. 6 A-C, aligned with the crRNA. The four Cmr4 cleavage sites are indicated on the sequences, and the respective Cmr4 subunits are shown (color coded as in fig 1. B-C)

(G) and (H) ssDNA and ssRNA cleavage time course experiments, respectively, in the presence and absence of ATP, comparing the wt CmrB-CTR complex (left panels) with the delta-7 mutant (right panels). The gels are one of 3 replicates used for the quantification plots in fig. 6E and 6F.
